# Dermomyotome-derived endothelial cells migrate to the dorsal aorta to support hematopoietic stem cell emergence

**DOI:** 10.1101/2020.05.14.096305

**Authors:** Pankaj Sahai-Hernandez, Claire Pouget, Ondřej Svoboda, David Traver

## Abstract

Development of the dorsal aorta is a key step in the establishment of the adult blood-forming system, since hematopoietic stem and progenitor cells (HSPCs) arise from ventral aortic endothelium in all vertebrate animals studied. Work in zebrafish has demonstrated that arterial and venous endothelial precursors arise from distinct subsets of lateral plate mesoderm. Earlier studies in the chick showed that paraxial mesoderm generates another subset of endothelial cells that incorporate into the dorsal aorta to replace HSPCs as they exit the aorta and enter circulation. Here we show that a similar process occurs in the zebrafish, where a population of endothelial precursors delaminates from the somitic dermomyotome to incorporate exclusively into the developing dorsal aorta. Whereas somite-derived endothelial cells (SDECs) lack hematopoietic potential, they act as local niche to support the emergence of HSPCs from neighboring hemogenic endothelium. Thus, at least three subsets of endothelial cells (ECs) contribute to the developing dorsal aorta: vascular ECs, hemogenic ECs, and SDECs. Taken together, our findings indicate that the distinct spatial origins of endothelial precursors dictate different cellular potentials within the developing dorsal aorta.

## Introduction

The primitive vascular network, which integrates into all organ systems in the developing organism, arises from endothelial precursors termed angioblasts (Risau and Flamme, 1995). To form a functional vascular network, angioblasts must first differentiate into a variety of distinct arterial and venous endothelial cell (EC) types, including hemogenic, endocardial, and blood brain barrier ECs (Aird, 2007; Herbert and Stainier, 2011). EC differentiation is thought to initiate midway through somitogenesis, as angioblasts migrate to the embryonic midline and coalesce to form the vascular tube (Herbert et al., 2009; Isogai et al., 2003; Jin et al., 2005). The current view states that equipotent angioblasts undergo successive steps of differentiation that, along with cues from local microenvironments, give rise to specialized subsets of ECs (Atkins et al., 2011; Marcelo et al., 2013). However, this view does not take into account putative differences in EC function as a result of different developmental origins. Rather, exposure to embryonic signaling cascades mediated via Wnt (Hubner et al., 2017), Hedgehog (Hh)(Gering and Patient, 2005; Vokes and McMahon, 2004; Wilkinson et al., 2012; Williams et al., 2010), Vascular Endothelial Growth Factor (VEGF) (Casie Chetty et al., 2017; Hong et al., 2006; Lawson et al., 2003; Lawson et al., 2002; Wythe et al., 2013), and Notch molecules are thought to differentially instruct equipotent angioblasts to each distinct endothelial cell fate (Fang et al., 2017; Lawson et al., 2001; Siekmann and Lawson, 2007; Zhong et al., 2001).

Lateral plate mesoderm (LPM) is known to be the primary source of ECs across vertebrate phyla (Potente and Makinen, 2017). However, recent findings have suggested that ECs can arise from distinct mesodermal derivatives, including extraembryonic-derived erythromyeloid progenitors (EMPs), that contribute extensively to the murine kidney vasculature (Plein et al., 2018). Similarly, there is evidence in rodents and other amniotes, that paraxial mesoderm (PM) also generates a contingent of ECs that contribute to the embryonic vasculature (Ambler et al., 2001; Esner et al., 2006; Noden, 1989; Pardanaud et al., 1996; Pouget et al., 2006; Pouget et al., 2008; Wilting et al., 1995; Yvernogeau et al., 2012). Specifically, these ECs arise from a transient somitic compartment known as the dermomyotome (Eichmann et al., 1993; Ema et al., 2006; Pouget et al., 2008; Yvernogeau et al., 2012), where skeletal hypaxial muscle progenitors (skMPs) reside (Tozer et al., 2007). These skMPs give rise to migrating myoblasts that incorporate into the ventral body wall and limb musculature (Schienda et al., 2006) The existence of somite-derived endothelial cells, and their function, in other vertebrate species, such as zebrafish, is incompletely understood.

Here, using a combination of molecular and genetic approaches, we identify a population of somite-derived endothelial cells (SDECs) in zebrafish that arises from the dermomyotome. SDECs contribute mainly to the anterior portion of the dorsal aorta. Within the somite, EC-fate acquisition occurs in a sequential manner, concomitant with the epithelialization of each somite and the migration of angioblasts towards the midline of the embryo. We show that Wnt signaling is a key regulator of the distribution of ECs within the somite, whereas Notch signaling is necessary for skeletal muscle progenitor cell maintenance. Finally, epistasis experiments indicate that SDECs arise from bipotent precursors within the somite, with skMPs showing competency to become ECs in a NPAS4l (cloche (Stainier et al., 1995)) –dependent manner. Single cell RNA sequencing (scRNA-seq) of purified ECs identified SDECs as well as additional subsets of EC types with distinct molecular signatures. Collectively, these findings shift the current paradigm of vascular origins and indicate that there are distinct pools of endothelial progenitors that are molecularly and functionally distinct as early as the end of gastrulation.

## Results

### Somites Give Rise to Etv2^+^ Endothelial Cells Concomitant with Somite Epithelization

In zebrafish, most ECs originate from the LPM (Jin et al., 2005); however, recent studies have suggested that somites also produce ECs that integrate into the vascular cord (Nguyen et al., 2014), but these remain incompletely defined. Therefore, we first aimed to characterize the development of SDECs. To accomplish this, we used confocal time-lapse imaging in *etv2:GFP* transgenic zebrafish (Veldman and Lin, 2012), injected with mOrange2:CAAX mRNA, which have early endothelial cells marked by GFP with cell boundaries demarcated with mOrange2. As previously described (Veldman and Lin, 2012), beginning at the 10 somite stage (ss), we observed a line of *Etv2:GFP*^*+*^ ECs along the most medial part of the LPM (Figure 1A); we did not detect any *Etv2:GFP*^*+*^ cells in the somite at this stage (Figure 1A). Initiating at the 12 ss, we noted *Etv2:GFP*^*+*^ cells in the lateral lip of the somitic compartment (Figure 1B). After the onset of Etv2:GFP expression, somite-derived Etv2:GFP+ cells round up (Figure 1C and 1D), delaminate from the somite, then integrate into the cohort of LPM-derived Etv2:GFP+ cells as they migrate towards the midline (Figure 1E, Supplementary Movie S1). These results indicate that there is a rare population of ECs that arise in sequential fashion starting at somite 12.

**Figure 1:**
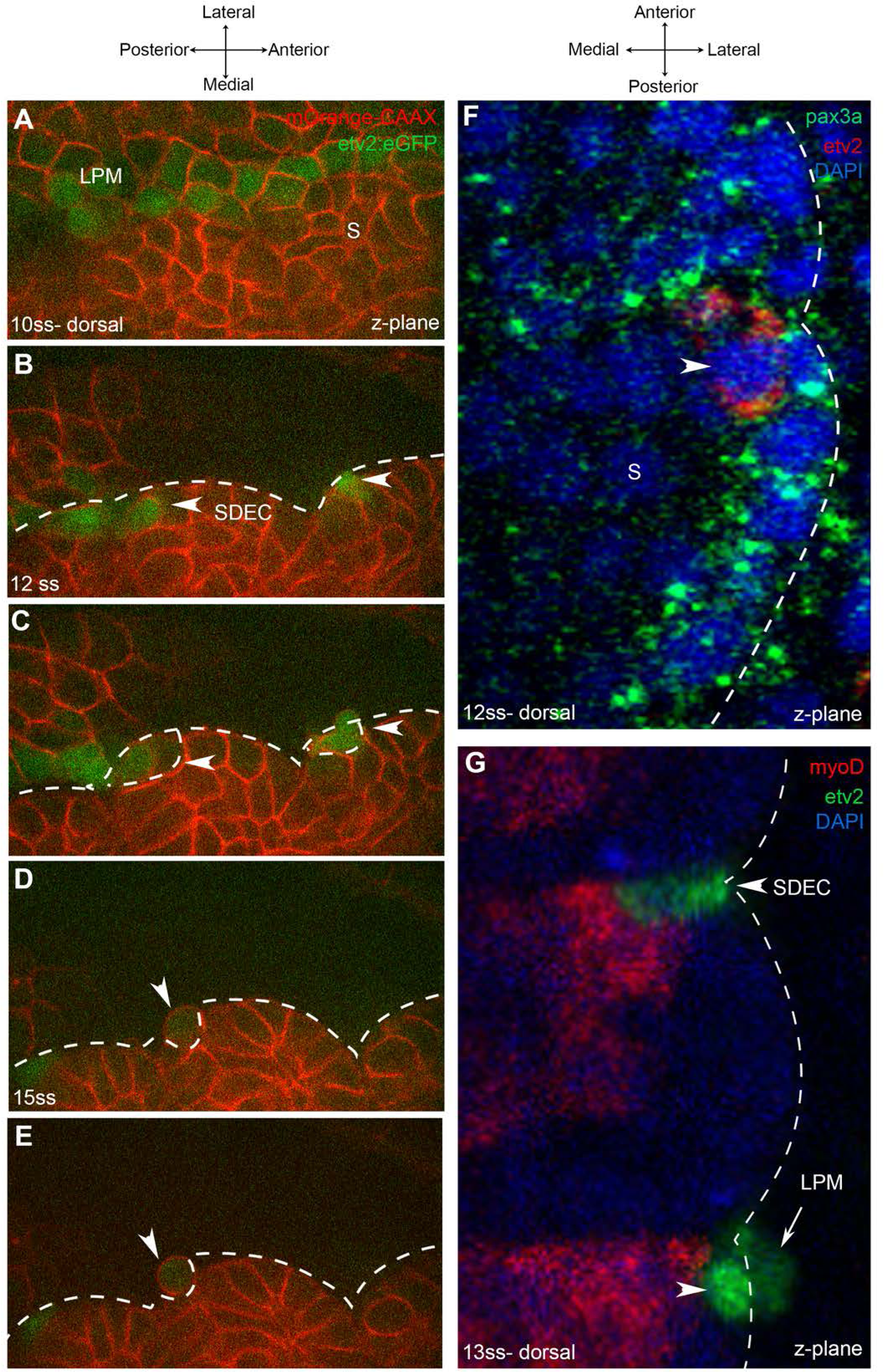
Endothelial cells emerge from the dermomyotome at 12ss. **A-E**. Time-lapse imaging from a dorsal view of *Tg(Etv2:GFP)* embryos injected with mOrange:CAAX mRNA and imaged between 10ss to 15ss. **A**. The expression of Etv2:GFP+ cells is visible along the LPM region. At this stage, no Etv2:GFP+ cells are visible in the somites. **B**. Starting at 12ss, the first Etv2:GFP^+^ cells are detected in the lateral lip of the dermomyotome (arrowheads). Simultaneously, the LPM Etv2:GFP^+^ cells start migrating to the midline. **C**. Soon after emergence, somite derived Etv2:GFP^+^ cells change shape and become rounder (arrowheads). **D-E**. Etv2:GFP^+^ cells bud off from the somite as individual cells (arrowhead). **F**. Dorsal view of a 12ss embryo submitted to a double fluorescent *in situ* hybridization for muscle progenitor maker, Pax3a (green), and endothelial marker, Etv2 (red). Pax3a expression reveals the dermomyotome compartment that contains the muscle progenitor cells. An Etv2+ cell (red) is found in the dermomyotome, co-expressing Pax3a (green and arrowhead), showing co-localization of an endothelial and muscle progenitor cell marker. **G**. Somitic Etv2+ cells (green) do not co-express the muscle differentiation marker myoD (red), suggesting that Etv2+ expression is restricted to the muscle progenitor region of the somite. Dashed white lines delimitate somite from the LPM (arrow).

In mouse and chicken, SDECs emerge from the same region as skeletal muscle progenitor cells, in the dermomyotome compartment (Tozer et al., 2007). To examine the spatial origins of SDECs in the zebrafish, we performed double fluorescent *in situ* hybridization (FISH) for the endothelial marker *etv2* and the skeletal muscle progenitor marker *pax3a* (Relaix et al., 2005) between 12-14ss, where we observed *etv2*^*+*^ cells to co-localize with *pax3a*^*+*^ cells in the somite (Figure 1F). However, FISH for *etv2* and *myod*, a marker of differentiated muscle cell types (reviewed in Hernandez-Hernandez et al., 2017), did not show co-localization (Figure 1G). Together, these results suggest that SDECs emerge within the somite from precursor cells shared with the muscle lineage. Because somitic *Etv2*^*+*^ cells are localized specifically within the dermomyotome region, the muscle progenitor cell compartment of the somite, a conserved mechanism of SDEC generation may be shared among vertebrates (Pouget et al., 2006; Mayeuf-Louchart et al., 2014; Mayeuf-Louchart et al., 2016).

### The Dermomyotome Contains Progenitors with Muscle and Endothelial Potential

Since our results above suggested the presence of bipotent progenitors with competence for muscle and endothelial cell differentiation, we sought to determine if blocking skeletal muscle differentiation may lead to enhanced SDEC generation by knocking down *mesenchyme homeobox 1* (*meox1)*, an important regulator of muscle cell formation (Mankoo et al., 2003). Using a *meox1* translation blocking morpholino (MO) (Nguyen et al., 2014) and FISH, we observed ectopic formation of *etv2, meox1* double positive cells in morphant animals (Figure 2A-F). Like normal SDECs, these ectopic SDECs began emerging between 12-16 ss; however, *meox1* morphants continued to generate SDECs as late as 26 ss (Figure 2G, H), which we have never observed in wild type animals. Therefore, Meox1 loss of function leads to enhanced and prolonged production of SDECs.

**Figure 2:**
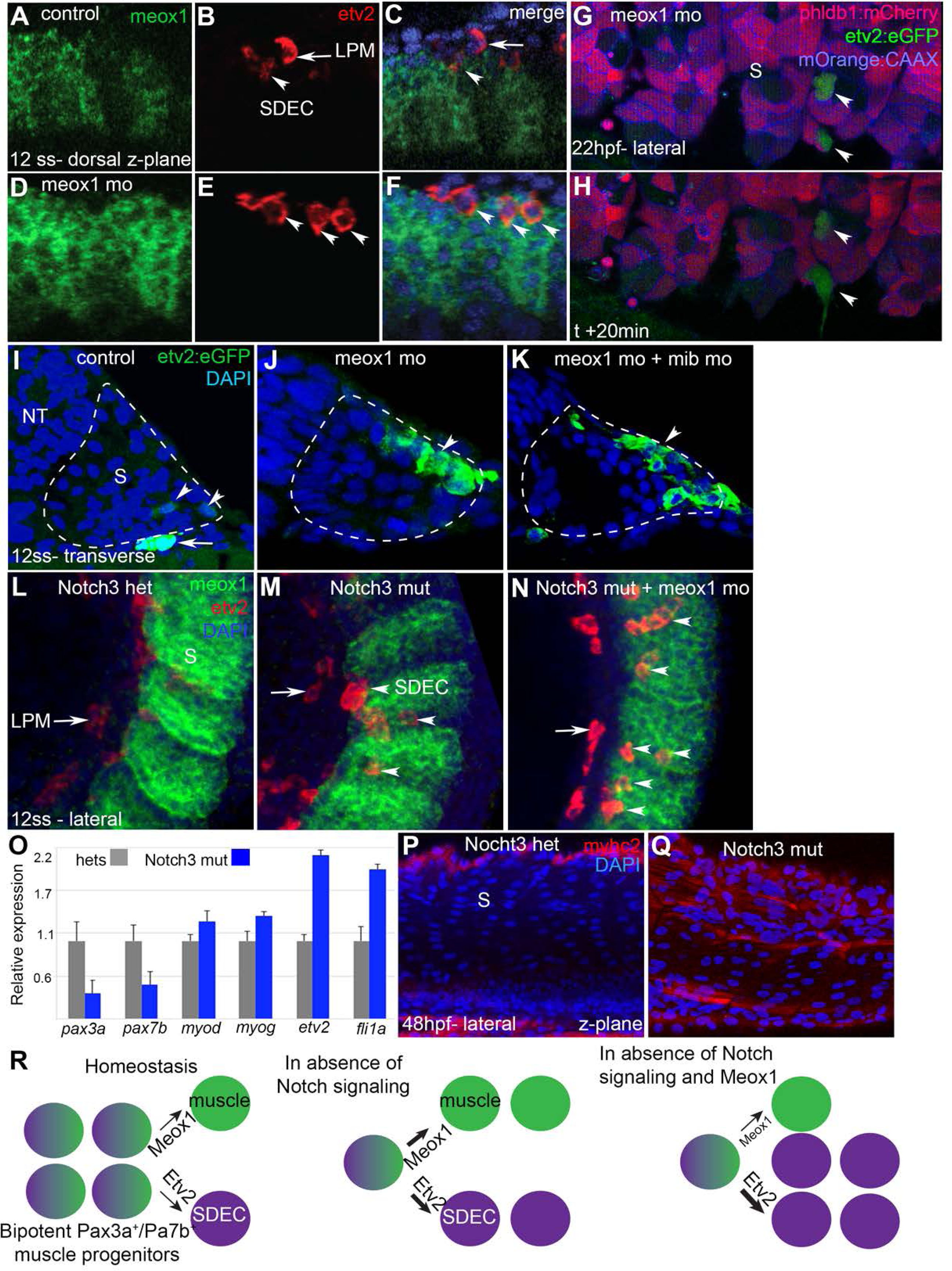
Notch is required for the maintenance of a bipotent skeletal muscle progenitor population in the somite. **A-F**. Dorsal view of 12ss control (A-C) and *meox1* morpholino (D-F) injected embryos. Embryos were submitted to double fluorescent *in situ* hybridization for Meox1 (green) and Etv2 (red). In control and morphant embryos, Meox1, Etv2 double positive cells are detected within the somite compartment (arrowhead). **C, F**. Overlay of Meox1 (green), Etv2 (red), and DAPI (blue). **D-F**. Knockdown of *meox1* results in ectopic formation of double positive cells within the somite (arrowhead). **G, H**, Time-lapse imaging of a 22hpf (26ss) *Tg(phldb1:Gal4:UAS-RFP; Etv2:GFP)* embryos, injected with *meox1* morpholino and mOrange2:CAAX mRNA, to delineate cell boundaries. Knockdown of *meox1* results in an extension of the time-period that the dermomyotome can generate Etv2:GFP^+^ cells (arrowheads). **I-K**, Cross section of 12 ss *Tg(Etv2:GFP)* control embryos. In absence of *meox1* (**J**), ectopic Etv2:GFP^+^ cells are visible in epithelialized layer of the somites, compared to controls (**I**). In embryos coinjected with *mib* and *meox1* morpholinos, the number of Etv2:GFP^+^ cells within the somite compartment (outlined compartment), is substantially increased **(K)**, suggesting that Notch signaling is dispensable for SDECs specification. **L-N**, Lateral view of 12ss embryos analyzed by FISH for *meox1* (green), *etv2*(red), and DAPI (blue). In Notch3 heterozygotes control **(L)** and mutant embryos **(M)**, SDEC etv2^+^ cells are detected in the somites. **N**. Notch3 mutant embryos co-injected with *meox1* morpholino, results in ectopic formation of Etv2 Meox1 double positive cells (arrowhead). **O**. qRT-PCR in 24hpf Notch3 mutant embryos and sibling controls. Genetic ablation of Notch3 results in decreased expression of muscle progenitor marker *pax3a* and *pax7b*; increased expression of muscle differentiation gene, *myod* and *myog* and endothelial markers, *etv2* and *fli1*. All genes analyzed between Notch3^-/-^ mutant and Notch3^-/-^ het embryos showed a statistically significant difference (p<0.001, Student’s t test; n=3.) **P**,**Q**, Notch3 mutant embryos show premature expression of MyoHII in 48hpf embryos **(Q)** compared to sibling controls **(P). R**. Summary cartoon for the role of Notch Signaling in the maintenance of bipotent-muscle progenitors (bipotent muscle progenitors in purple and green; muscle cells in green; endothelial cells in purple).

Previous work in mouse identified the Notch signaling pathway as a positive regulator of EC over muscle fate in the somite (Mayeuf-Louchart et al., 2014). We tested the requirement for Notch signaling in zebrafish SDEC generation by knocking down the essential Notch regulator *mindbomb (mib)* (Itoh et al., 2003) in *Etv2:GFP*^*+*^ animals and examining SDEC formation between 12-14 ss. While loss of Meox1 alone led to ectopic formation of a few SDECs (Figure 2I, J), double knockdown of *mib* and *meox1* led to a profound expansion of *Etv2:GFP*^*+*^ cells within the somite (Figure 2K). These results led us to hypothesize that Notch signaling is dispensable for *Etv2:GFP*^*+*^ cell formation but required for skeletal muscle progenitor maintenance. To examine the role of Notch signaling more precisely in these cells, next we queried *Notch3*^*-/-*^ animals, since it is the primary Notch receptor in the somites at this stage (Kim et al., 2014). We injected *Meox1* MO into *Notch3*^*-/-*^ embryos and performed FISH for *meox1* and *etv2* (Figure 2L-N). Using this strategy, we observed that in *Notch3*^*-/-*^ homozygous mutant embryos, the formation of SDECs was not impaired (Figure 2L, M). Moreover, following combined loss of Notch3 and Meox1, we observed the ectopic formation of *etv2*^+^, *meox1*^+^ double positive cells by FISH (Figure 2N and Supplementary Figure 1). Together, these results show that Notch signaling is dispensable for the specification of SDECs.

To determine which cell populations were specifically affected by loss of Notch3, we assessed the expression levels of markers for muscle progenitors (*pax3a, pax7b*), differentiated muscle cells (*myod, myog*) and differentiated endothelial cells (*evt2, fli1a*) by qRT-PCR in *Notch3*^*-/-*^ embryos at 24hpf (Figure 2O). In *Notch3*^*-/-*^ animals, we observed a decrease in expression of the muscle progenitor markers, concomitant with an increase in the expression of muscle differentiation markers and endothelial differentiation markers (Figure 2O). Furthermore, we observed premature expression of the muscle differentiation marker MyoHII (Sjoblom et al., 2008; Beier et al., 2011; Salucci et al., 2015) by antibody staining at 48 hpf in *Notch3*^*-/-*^ animals (Figure 2P, Q). Together, these results indicate that Notch3 signaling is required for Pax3^+^, Pax7^+^ muscle progenitor maintenance, and its absence leads to premature differentiation of muscle and SDEC fates (Figure 2R).

### NPAS4l is Required for the Formation of SDECs

*NPAS4l* (*cloche*) is regarded as the most upstream gene required for blood and EC specification (Stainier et al., 1995, Reischauer et al., 2016). We therefore sought to determine if *NPAS4l* is required for the development of SDECs. WISH at 12 ss embryos showed a complete absence of *etv2* expression along the embryonic A-P axis in *cloche* mutants compared to controls (Figure 3A-B), showing that *NPAS4l* function is required for both PLM-derived ECs and SDECs. As above, we knocked down the function of *meox1* in order to increase SDEC differentiation from bipotent somitic progenitors. SDECs were not observed in this scenario either, indicating NPAS4l function is necessary to generate the SDEC lineage (Figure 3C-D). To further increase the number of SDECs, we injected embryos from *Etv2:GFP*^*+/+*^; *cloche*^*-/+*^ adult pairs with *mib* Mo and *meox1* MO. Similarly, we observed a complete absence of *Etv2:GFP*^*+*^ cells in *cloche*^-/-^ mutant embryos, compared to *cloche* control siblings (*cloche*^-/+,^, *cloche*^+/+^) Figure 3E-H. Moreover, we performed FISH for *meox1* and *etv2* and did not observe any double positive cells in *cloche*^-/-^ mutants, compared to *cloche* control embryos (Figure 3I-N). Lastly, we performed qRT-PCR for endothelial and muscle cell genes from 48 hpf *cloche*^*-/-*^ mutant embryos compared to control *cloche* control embryos (Figure 3O). As expected, there was a decrease in the expression of the endothelial genes *etv2* and *fli1a*. Interestingly, however, we also observed a concomitant increase in expression of skeletal muscle differentiation genes *myod* and *myogenin* (Figure 3O). Together, these results confirm that *NPAS4l* (*cloche*) is required for the specification of endothelial cells from the PLM and establish a similar requirement for SDEC generation from shared skeletal muscle progenitor cells (Figure 3P, Q).

**Figure 3:**
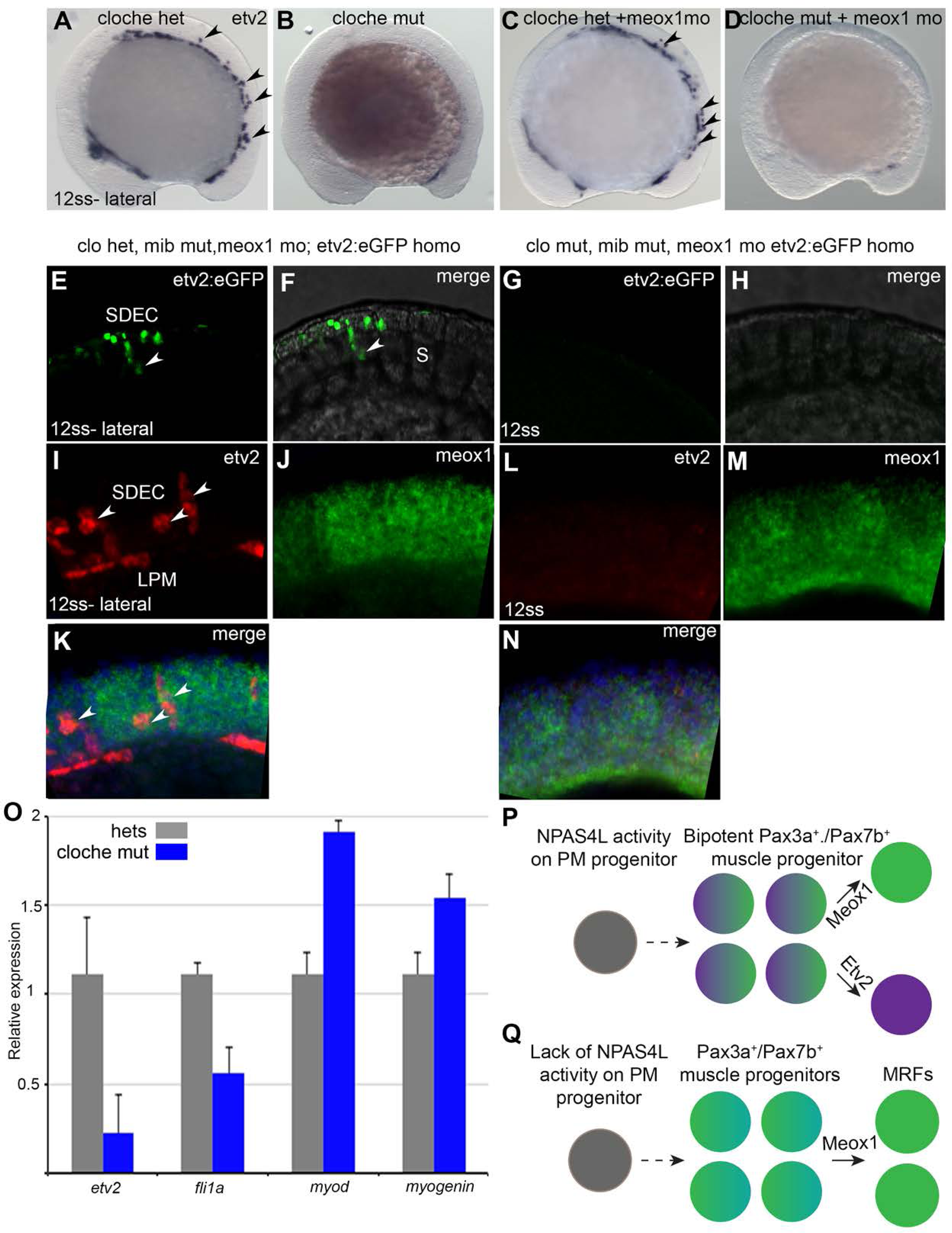
NPAS4l is required for the specification of the SDECs. **A-D**. WISH for etv2 in 12ss embryos. **B**. *cloche* mutant embryos shows absence of etv2 expression along the A-P axis of the embryo, compared to sibling control **(A). D**. Similarly, cloche mutant embryos injected with Meox1 morpholino, results in loss of etv2 expression, compared to sibling control **(C). E-H**. Similarly, *Tg(Etv2:GFP*^*+/+*^*(homozygous); cloche*^*-/+*^) adult fish were incrossed and embryos were injected with both meox1 and mib morpholino. **G, H**. Cloche mutant embryos show loss of etv2:GFP expression in the LPM and somites at 12ss compared with control embryos (**E, F** arrowheads). **I-N**. *Tg(cloche*^*-/+*^) adult fish were incrossed and embryos co-injected with mib and meox1 morpholino, and performed a FISH for meox1 (green) and etv2 (red) at 12ss. **L-N**. cloche mutants shows loss of all etv2, meox1 double positive cells. **O**. qRT-PCR of cloche mutant embryos shows the expected loss of endothelial genes and concomitant increase of muscle differentiation genes. All genes analyzed between Cloche^-/-^ mutant and Cloche^-/-^ het embryos showed a statistically significant difference (p<0.001, Student’s t test; n=3.) **P, Q**. Summary model by how NPAS4l might affect endothelial cell competence in PM progenitors (early mesoderm progenitor in grey; bipotent muscle progenitor in purple and green; muscle cells in green; endothelial cells in purple).

### Wnt Signaling Regionalizes the Formation of SDECs

Previous work has shown that Wnt signaling is required for the differentiation of muscle cells, through activation of the required skeletal muscle factor Myf5 reviewed in ((von Maltzahn et al., 2012)). In addition, inhibition of Wnt signaling in early PM leads to an increase in endothelial cells that can integrate into the zebrafish vasculature (Veldman et al., 2013), suggesting that Wnt signaling may also act later in the somite to balance muscle and endothelial cell production from shared muscle progenitor cells. To confirm that Wnt signaling is active in *meox1*^*+*^ muscle progenitor cells during SDEC development, we performed FISH for *meox1* in the background of a destabilized Wnt/TCF GFP-reporter line (Moro et al., 2012). We observed cells positive for both GFP and *meox1* (Figure 4A), indicating that Wnt signaling is active while SDEC fate decisions are occurring. Next, to determine if Wnt inhibition affects SDEC development, we treated *etv2:GFP*^*+*^ embryos with IWP-L6, a potent inhibitor of Wnt protein secretion (Wang et al., 2013), from 2-12 ss. Inhibition of the Wnt signal was confirmed by qRT-PCR for the canonical target gene *axin2* (Figure 4B), which led to a decrease in *meox1* and an increase in *etv2* transcripts. By examining serial sections, Wnt inhibition led to the formation of ectopic *Etv2:GFP*^*+*^ cells in the somites (Figure 4C, D), which we confirmed by performing time-lapse imaging of the same experimental setup between 10 ss and 14 ss (Supplementary Movie S2). Finally, we also confirmed that Wnt signaling is required for *meox1* expression (Figure 4E-F’). Together, these results show that Wnt signaling regionalizes the formation of SDECs within the somite within the most lateral region, and that its inhibition results in the ectopic formation of *Etv2:GFP*^*+*^ SDECs.

**Figure 4:**
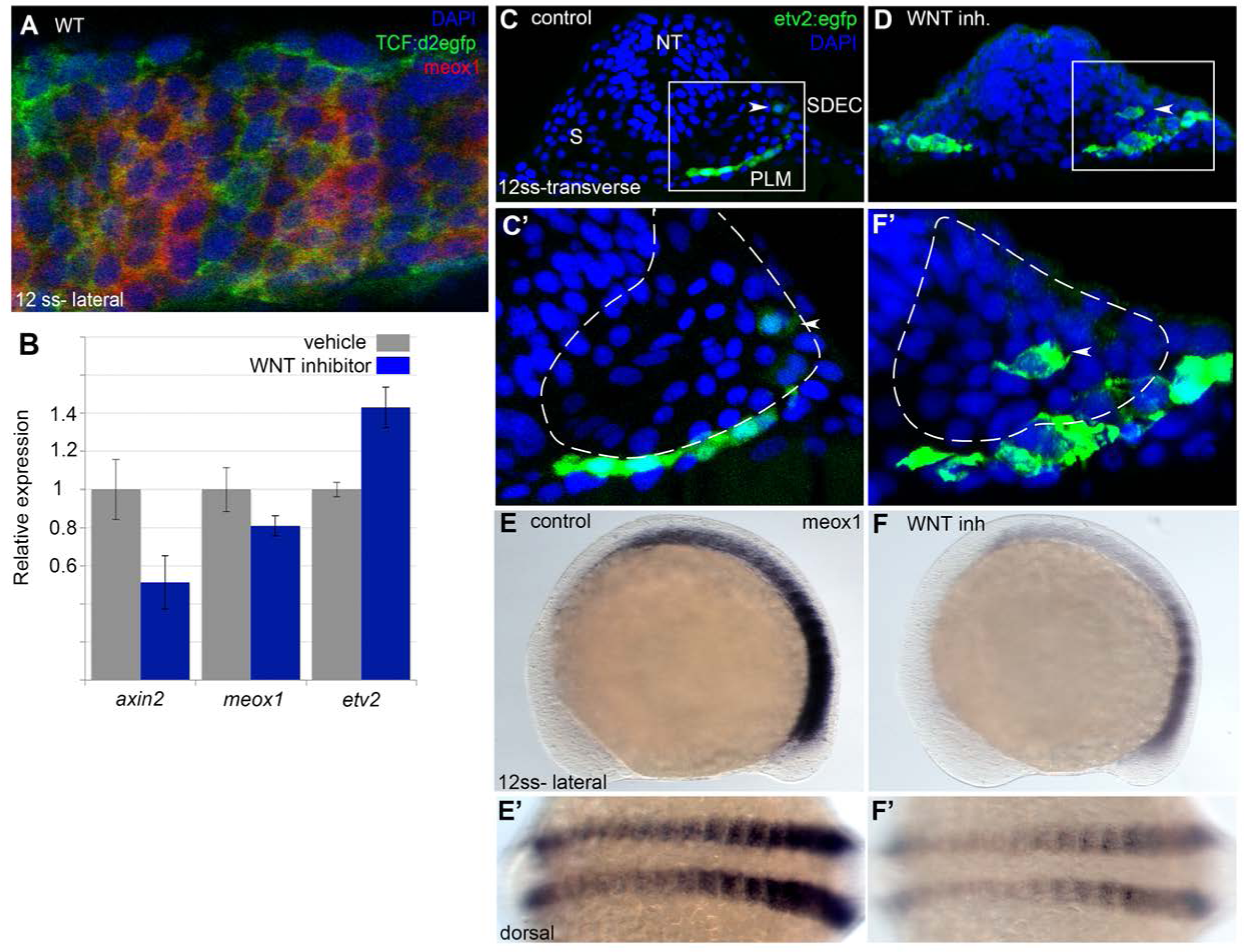
Wnt signaling is required for the regionalization of SDECs. **A**. FISH for meox1 (red) and antibody staining for a destabilized Wnt/TCF reporter line (green), shows somitic cells co-expression of GFP and Meox1 within the somite compartment. **B**. Inhibition of Wnt signaling using the chemical inhibitor IWP2 from 2ss to 15ss stage, results in decrease expression by qRT-PCR of axin2, meox1 with a concomitant increase of the expression of etv2. All genes analyzed between Wnt inhibitor and control embryos showed a statistically significant difference (p<0.001, Student’s t test; n=3.) **C, D**. Cross section of *Tg(Etv2:GFP)* embryos treated with IWP2 from 2ss-15ss stage, results in ectopic formation of Etv2:GFP^+^ cells within the somite, **D** (arrowhead). **E, F**. IWP2 treated embryos from 2ss-12ss stage results in decrease expression of Meox1 by WISH, compared to control embryos.

### SDECs Integrate into the Dorsal Aorta but do not Generate HSPCs

Next, we were interested in understanding the contribution of SDECs to the developing zebrafish vasculature. To this end we obtained a PM-specific Gal4 transgenic line, *Tbx6:Gal4*, which has been shown to recapitulate *tbx6* mRNA expression (Yabe et al., 2016). We crossed this line to *UAS:CRE*, which expresses the Cre recombinase upon Gal4 induction only within the PM, and to a ubiquitously expressed reporter line *βActin:>BFP>dsRed*, which upon genetic recombination switches from a BFP to dsRed expression cassette (abbreviated as A2BD) for clarity. Upon confocal imaging of the triple transgenic *tbx6:Gal4; UAS:Cre; A2BD* animals at 48hpf, we observed dsRed+ cells within the region of the axial vasculature (Figure 5A-C). To further confirm that these cells were ECs that arise from a Tbx6+ somitic population, we crossed the *Tbx6:Gal4; UAS:Cre* line with a previously published vascular-specific switch line, *Kdrl:CSY* (Zhou et al., 2011) The latter transgene is driven by an endothelial-specific promoter, and upon Cre-induced recombination switches ECs from CFP to YFP expression. When we imaged the different portions of the DA (Figure 5D) in *Tbx6:Gal4; UAS:Cre; Kdrl:CSY* triple transgenic embryos via confocal microscopy at 4 days post fertilization (dpf), we observed YFP+ cells that localized preferentially to the anterior region of the dorsal aorta (Figure 5E-G, K, Supplementary Figure 2). Notably, the most anterior region of the DA contained a higher percentage of YFP+ cells (Figure 5E, K), with the contribution of YFP+ cells decreasing in an anterior to posterior manner (Figure 5G, C). We found this labeling pattern to be consistent among different embryos over a range of developmental time points.

**Figure 5:**
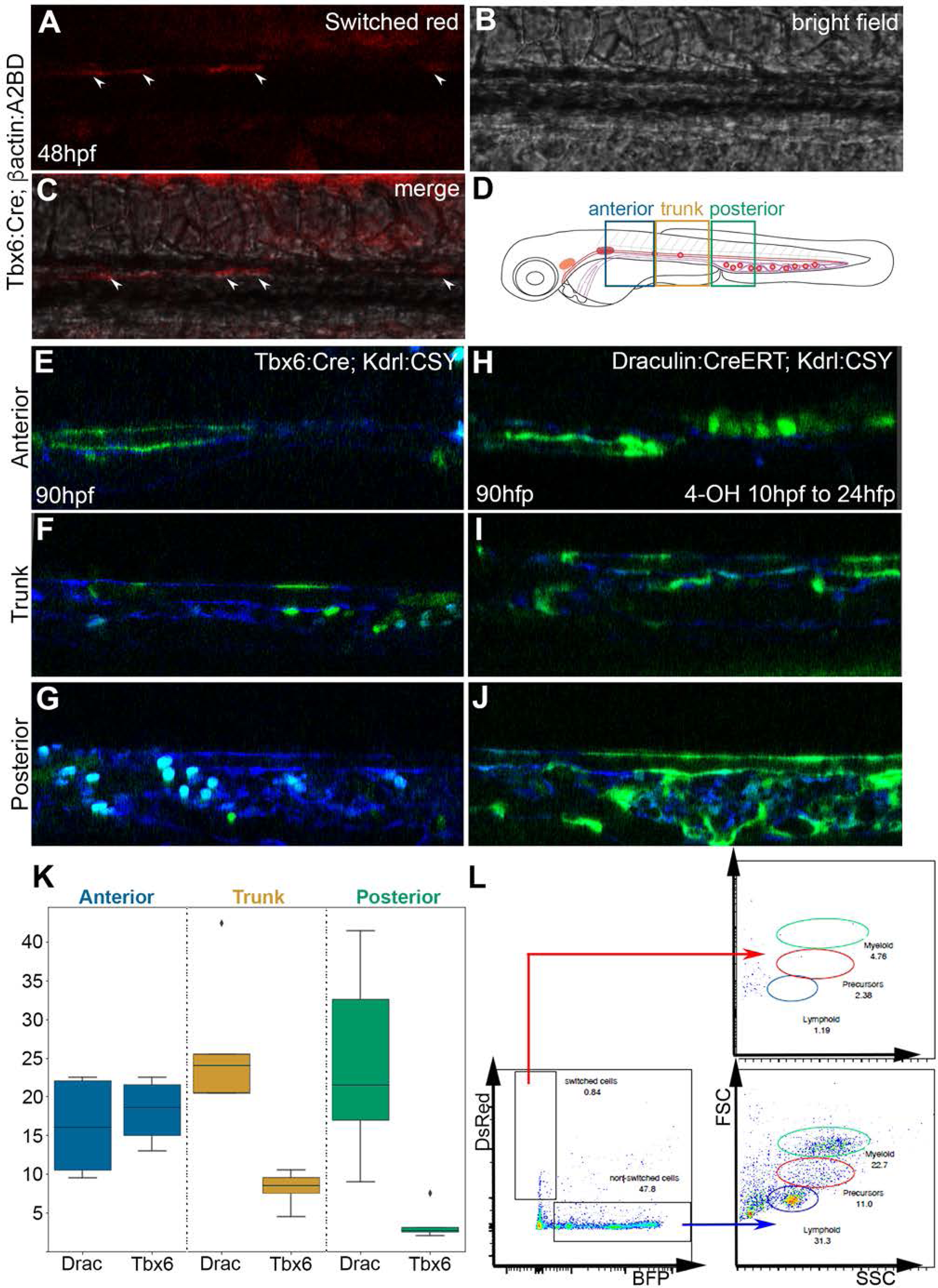
SDECs contribute to the dorsal aorta and do not generate HSPCs. **A-C**. Lineage tracing of somite derived cells using *Tg(Tbx6:Gal4; UAS:CRE; beta-actin:A2BD)* shows dsRed+ cells in the vasculature region at 48hpf (arrowheads). **E-J**. Using a vasculature specific switched line *Tg(Kdrl:CSY)*, we observe the contribution of SDECs or PLM-derived endothelial cells to the vasculature. **E-G**. For SDECs labelling, a PM-specific driver *Tg(Tbx6:Gal4;UAS:CRE)* was used. PM-derived YFP^+^ cells are observed in the vasculature of imaged embryos. **H-J**. For LPM-specific labeling, a *Tg(Draculin:CreERT2)* was used and treated with 10um of tamoxifen starting at 8 hpf. YFP^+^ cells are observed in all region of the vasculature. **K**. Quantification of YFP^+^ endothelial cells from Tbx6 or Draculin switched embryos. These quantifications where based from independent experiments per transgenic background with n=23 for Tbx6 switched embryos and n=9 for Draculin switch embryos. **L**. Analysis of the adult kidney marrow of *Tg(Tbx6:Gal4; UAS:CRE; beta-actin:A2BD)* shows no contribution to definitive blood cells (DsRed) through flow cytometry analysis whereas the FSC/SSC distribution of the unswitched BFP^+^ cells correspond to all blood lineages (quantifications based from independent experiments with a total of n=21 samples).

To complement these results, we performed lineage tracing using a LPM-specific switch line, *Draculin-CreERT2* (Henninger et al., 2017), likewise crossing it to *Kdrl:CSY* for EC-specific reporting. We incubated embryos with tamoxifen (4-OHT) to induce Cre-based recombination between 8 and 24 hpf. Confocal imaging showed complementary results to those from our PM-specific Tbx6+ experiments (Figure 5H, J). We observed that Draculin-derived YFP+ cells contribute to all regions of the vasculature, as would be expected from an LPM source. However, we observed the contribution of these cells was more robust within the posterior region of the DA compared with the Tbx6-labelled SDECs, consistent with our PM lineage tracing data (Figure 5E, K). Together, these results demonstrate that SDECs integrate into the DA in zebrafish and appear to contribute preferentially to the anterior portion of the dorsal aorta.

Since HSPCs derive from hemogenic endothelium specifically within the DA, we examined whether or not SDECs can generate HSPCs. We lineage traced PM-derived cells into the kidney, the adult hematopoietic organ in teleosts. We dissected kidneys of adult *Tbx6:Gal4; UAS:Cre; A2BD* transgenic animals and observed no contribution of switched dsRed+ cells to any hematopoietic lineages at this stage (Figure 5D). As a positive control for hemogenic endothelium, a ubiquitous vascular-specific transgenic Cre driver was utilized, *Kdrl-Cre; A2BD* (Supplementary Figure 3). Taken together, these results demonstrate that SDECs integrate into the dorsal aorta but do not generate HSPCs.

### Molecular Differences in ECs Foster Cellular Diversity within the Vasculature

The discovery of SDECs in zebrafish prompted us to revisit EC diversity in the developing zebrafish embryo using single cell RNA sequencing (scRNAseq). To do so, we collected EC enriched samples from distinct vascular transgenic animals, with the idea of representing diverse endothelial cohorts. Specifically, we purified cells via fluorescence-activated cell sorting (FACS) from *Etv2:kaede, Fli1:dsRed; Tp1:GFP*, and *Draculin:Dendra:H2B* between 22-24 hpf from synchronized embryos and we performed scRNAseq on each sample. Following quality control (QC) and basic clustering of each sample, we merged individual datasets. Altogether, we obtained a total of 10,160 single cells, 2,996 cells of which belonged to the endothelial lineage. Via unsupervised clustering of single cell transcriptomes and based upon known lineage associations, we identified six distinct endothelial cells clusters (Figure 6A). Using known genes within established lineages, we named each cluster based on likely tissue origins (Supplementary Sheet1). First, we identified an endothelial cell cluster 1 that contained the expression of paraxial mesoderm (PM) signatures genes, including *fras1, fn1b, fbn2b, acta1a* and *fgf8a*. This PM signature suggested this population to represent SDECs. Cluster 2 (Endocardium) likely represents endocardial ECs based upon co-expression of canonical heart genes, including *gata5, hand2* and *tbx20*. Cluster 3 (Kidney vascular endothelium cells, KVECs) co-expressed kidney-associated genes, such as *pax2a, pax2b, jag2b*, and *osr1*. Cluster 4 (Brain vascular endothelium cells, BVECs) co-expressed brain-associated genes, including *tbx*1 and *eya1*. Finally, we identified a large endothelial cluster 5 (General endothelium) that co-expressed canonical endothelial genes as well as a likely hemogenic cluster directly adjacent to it that co-expressed endothelial and hematopoietic genes, including as *gfi1aa, myb* and *cebpa*. Therefore, we merged clusters 6-7 that were adjacent to the General endothelium cluster, and we termed it hemogenic endothelium. Differential expression analysis among clusters identified distinct gene programs enriched within specific subsets (Figure 6B, Supplementary Sheet1). Thus, this scRNA-seq approach served to not only identify SDECs and hemogenic endothelium, but also a variety of additional tissue-specific EC subsets in the 22-24 hpf embryo.

**Figure 6:**
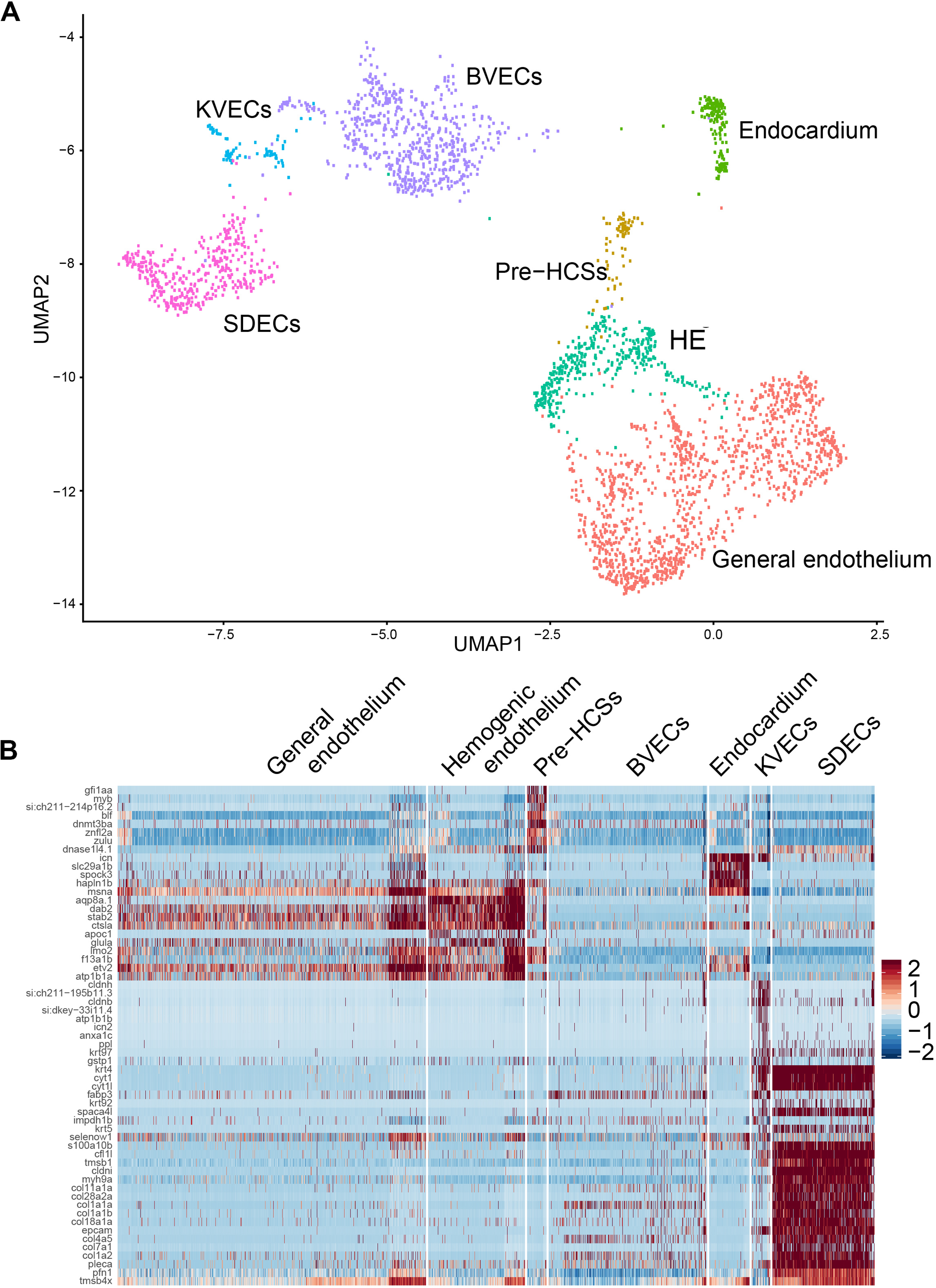
Cell clustering and cell-type-specific endothelial cell markers. **A**. Uniform Manifold Approximation Projection (UMAP) plots of 22hpf scRNA-seq data, 2,996 total cells that represent the endothelial lineage. Clusters were named according to their gene expression: somite derived endothelial cells (SDECs), Brain Vascular Endothelium (BVECs), Kidney Vascular Endothelial cells (KVECs), General Endothelium, Pre-HSCs Hemogenic Endothelium (HE), and Endocardium. Color-coded marker gene expression levels are shown on corresponding clusters. **B**. An expression heatmap of 22hpf single cell transcriptome showing the top predicted differentially expressed marker genes across the different clusters.

### SDECs Support the Emergence of HSPCs

Differential expression analysis of SDECs versus other endothelial clusters identified many genes previously attributed to ‘niche’ functions required for the induction of aortic hemogenic endothelium (Figure 7A) (Charbord et al., 2014). Notable among these is BMP4, which is required for the induction and maintenance of Runx1 expression within the hemogenic endothelium (Wilkinson et al., 2009). These results suggested that SDECs might be acting in a paracrine manner to support hematopoietic induction. If so, we reasoned that changes in the number of SDECs may result in altered formation of hematopoietic cells. Increasing the numbers of SDECs via enforced expression of *etv2* with a muscle-specific *mylz2* promoter (Ju et al., 2003), or knockdown of *meox1*, led to an increase in *runx1*^*+*^ expression by WISH (Figure 7B-E). By contrast, decreasing the number of SDECs via over expression of *meox1* mRNA led to the loss of *runx1*^*+*^ and *gata2b*^*+*^ hematopoietic cells in the DA (Figure 7F-I). Next, since *bmp4* expression appeared to be differentially regulated between SDECs and ECs of the PLM (Figure7A), we queried if its expression would be increased upon increase generation of SDECs. Upon loss of *meox1* function, we found that *bmp4* expression was increased by WISH at 24hpf in the DA (Figure 7J, K). Taken together, these results demonstrate that SDECs act in a paracrine manner through BMP signaling to support the induction of hematopoietic cells from neighboring hemogenic endothelial cells.

**Figure 7:**
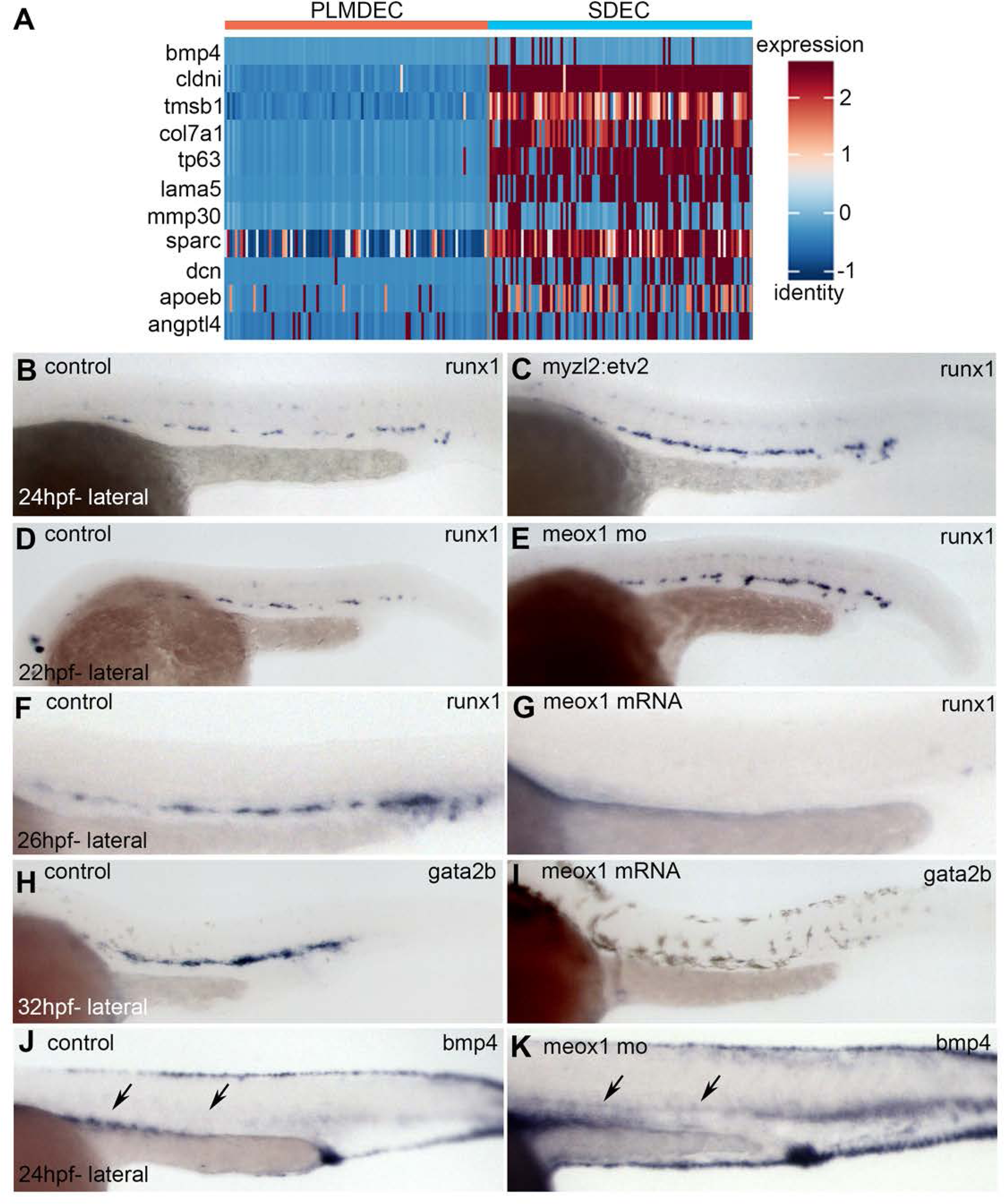
SDECs act as a vascular niche population for the hemogenic endothelium. **A**. Heatmap of genes that are differentially regulated between endothelial cells of the PLM and SDECs. **B, C**. Zebrafish embryos were injected with a transgene containing a somite specific promoter, Myzl2, and driving an Etv2 transgene, to ectopically induced SDECs. By WISH we observe an increase of *runx1* expression of 24hpf injected embryos, compared to embryos injected with an empty Myzl2 vector. **D-E** Similarly, Meox1 morphant embryos exhibit an increase expression of runx1 by WISH, compared to uninjected control embryos. **F-I**. Conversely, overexpression of Meox1 by mRNA results in a strong reduction of the hemogenic markers *runx1* and *gata2b*. **J-K**, Knockdown of meox1 results in increase expression of bmp4 in the dorsal aorta region by WISH.

### Early Notch Signaling Regulates the Rate of Differentiation of Endothelial Cells

Our results thus far have shown that SDECs integrate exclusively to the dorsal aorta, act in a paracrine manner to support the formation of hematopoietic cells, and do not give rise to HSPCs. With the noted importance of Notch signaling in establishment of both arterial and hemogenic endothelium, we were interested in exploring how Notch signaling might affect the formation of SDECs, arterial cells and hematopoietic cells at the single cell level. Previous work has shown that global inhibition of Notch signaling leads to loss of arterial cell fate and hemogenic endothelium (Lawson et al., 2001) (Burns et al., 2005). Conversely, Notch overexpression results in the expansion of the hematopoietic program (Guiu et al., 2013). The exact molecular changes that occur following Notch modulation between hemogenic and arterial endothelium are difficult to assess since each population is closely related and the number of cells in the developing embryo are limited. We thus utilized scRNAseq to profile single *Etv2:Kaede*^*+*^ cells from 22 hpf *mib*^-/-^ or control (*mib*^-/+^ and *mib*^+/+^) sibling mutant embryos, which have global loss of Notch signaling. From control siblings, we FACS purified 2,817 single cells, 1,392 of which belong to the endothelial lineage. Similarly, in the Notch mutant, we purified 2,720 single cells, 1,008 cells of which belong to the endothelial lineage. Comparison of Notch mutants to control sibling cells identified 155 genes to be differentially expressed (Supplementary Sheet2). Figure 8A shows a heatmap of the 20 most upregulated and downregulated genes between mib mutant and control cells. Interestingly, arterial genes in *Etv2:kaede*^*+*^, *mib*^*-/-*^ mutant cells were expressed at 22 hpf (Figure 8H). This result was unexpected, since Notch signaling has been thought to be required for the establishment of the arterial cell program. Because mRNA abundance in RNAseq data only captures a static snapshot at a specific point in time, we decided to utilize RNA velocity techniques (Zywitza et al., 2018) to gauge the differentiation states of *mib*^-/-^ or control ECs.

**Figure 8:**
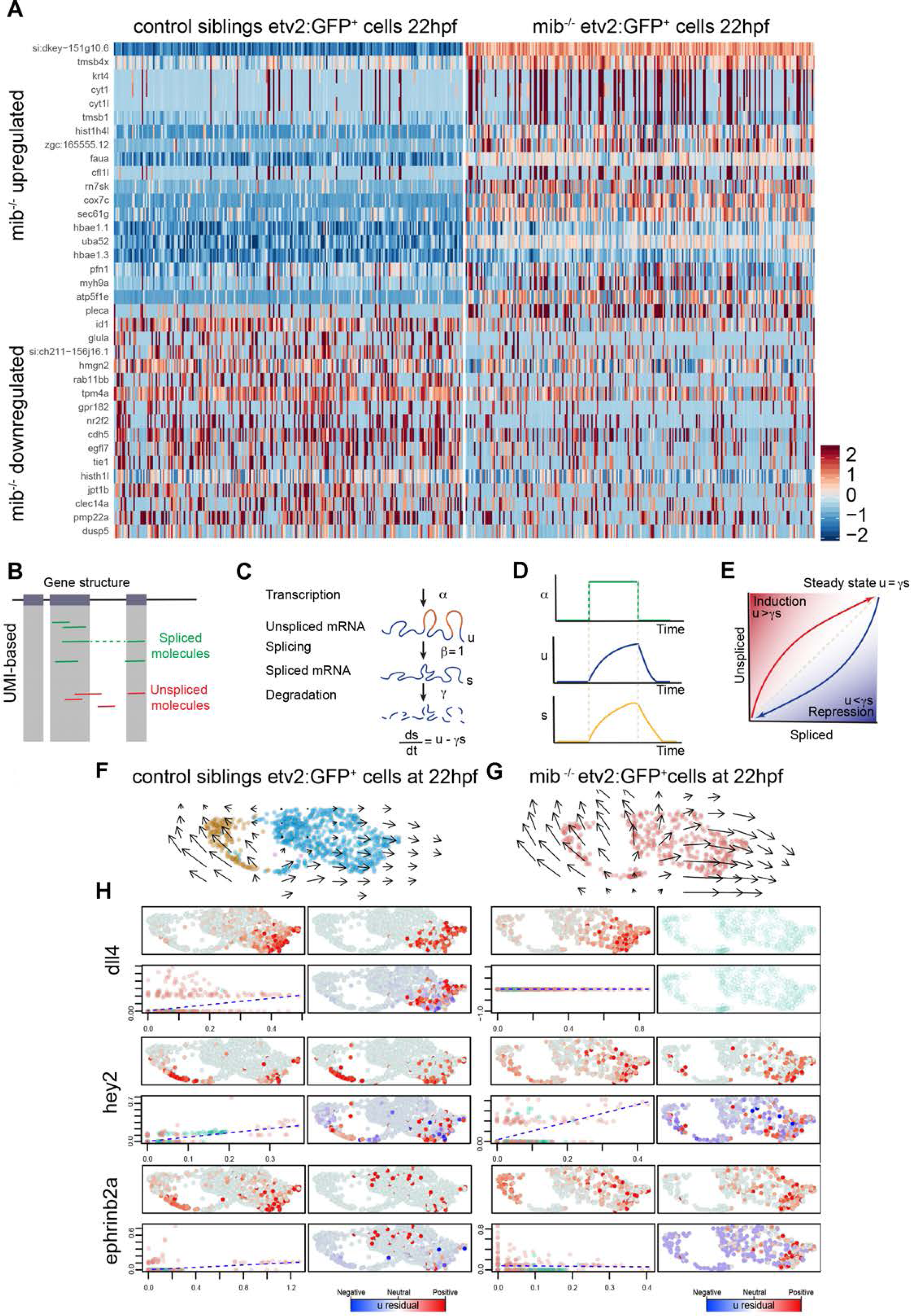
Early Notch signaling regulates the rate of differentiation of endothelial cells. **A**. Heatmap of the 20 most upregulated and downregulated genes between 1,392 mib^-/-^; and 2,817 control endothelial cells. **B-E**. Adapted from Zywitza et al., **B**. Spliced and unspliced counts are estimated by separated counting reads that incorporate intronic sequences. **C-E**. Model of transcriptional dynamics capturing transcription (alpha), splicing (beta), and degradation (gamma), rates involved in production of unspliced (u) and spliced (s) mRNA products and solution to the models. **F, G**. Velocities are visualized on a UMAP plot between Notch mutant **(G)** and control endothelial cells **(F)**, using Gaussian smoothing. **H**. Expression pattern of unspliced-spliced phase portraits, and residuals are shown for arterial genes, dll4, hey2 and ephrb2a between mib mutants (right panel) and control endothelial cell types (left panel).

RNA velocity uses the time derivative of gene expression state by estimating spliced versus unspliced mRNA from common RNA sequencing protocols. As an output, it generates high dimensional vectors that predict the future state of individual cells on a timescale of hours. During a dynamic process, such as a cell entering a differentiation program, an increase in the transcription rate alpha results in a rapid increase of unspliced mRNA, which is then followed by an increase in spliced mRNA, until a new steady state is reached (Figure 8B-E). Using RNA Velocity, we generated high dimensional vectors of *Etv2:Kaede*^*+*^ EC cell states (Figure 8F, G), as well as individual phase portraits (Figure 8H) for arterial genes. These results show that the induction of endothelial differentiation genes follow a pattern across the endothelial cell manifold. For example, we observe the arterial transcripts *dll4, hey2* and *ephb2a* to be induced at the periphery of ECs (cells with longer arrows) and are thus farther away in the differentiation hierarchy from ECs located within the medial portion of the cluster (shorter arrows) (Figure 8F).

To determine the consequences in the absence of Notch signaling in *Etv2:Kaede*^*+*^, *mib*^*-/-*^ ECs, we performed similar RNA velocity analysis. First, we observed that the directional flow of general ECs in Notch mutant endothelium to be more dynamic (Figure 8G; longer arrows) than control ECs (Figure 8F). These data suggest that Notch mutant ECs are further along the differentiation hierarchy than control *Etv2:Kaede*^*+*^ ECs. Upon closer inspection of the arterial differentiation gene program, we observe that arterial genes are prematurely expressed in Notch mutant *mib*^-/-^ when compared to control *Etv2:Kaede*^*+*^ ECs (Figure 8H). By comparing spliced and unspliced arterial genes, we noticed an expression hierarchy of *dll4, hey2* and *efnb2a*, respectively (Figure 8H). These results show that early Notch signaling is dispensable for the *initiation* of the aortic program but is required for the regulation of endothelial precursors entering its differentiation program. Thus, the early role of Notch signaling may be to maintain EC precursors in a plastic, or primed state.

Next, we were interested in assessing the difference in the hemogenic endothelium and pre-HSPC clusters of controls versus *mib*^-/-^; *Etv2:Kaede*^*+*^ cells. We computationally assigned control versus *mib*^-/-^; *Etv2:Kaede*^*+*^ cells to their respective, previously identified clusters. We observed that hemogenic-like cells (hemogenic endothelium and pre-HSPCs) still mapped within the population of *mib*^-/-^; *Etv2:Kaede*^*+*^ cells. However, we noticed that their numbers were significantly reduced in Notch mutant ECs, 166 cells in control vs 77 cells in *mib*^-/-^. In addition, specific comparison of the Pre-HSPC clusters showed that genes important for hematopoietic cell formation such as *lmo2* and *gpr182* were reduced in *mib*^-/-^ mutant cells (Supplementary Figure 4). Lastly, we compared the gene expression of SDECs between control versus *mib*^-/-^ *Etv2:Kaede*^*+*^ cells and did not observe a significant change in the SDEC gene program (Supplementary Figure 5). Together, these results suggest that Notch signaling plays a more central role in the specification of hemogenic endothelium rather than in the arterial or SDEC program itself.

### Inhibitor of Differentiation 1 (ID1) Works in Tandem with Notch Signaling to Enable the Specification of the Hemogenic Endothelium

One of the most highly differentially expressed genes arising from the Notch mutant single-cell analysis was Inhibitor of Differentiation 1 (ID1) (Figure 9A). As noted above, the RNA velocity data indicated that Notch signaling may serve as a gatekeeper for differentiation onset of ECs. Previous work has shown that Inhibitor of differentiation genes can maintain progenitors in an undifferentiated state and help facilitate choices between distinct cellular fates (Zhang et al., 2014) (Malaguti et al., 2019) (Leeanansaksiri et al., 2005) (Bedford et al., 2005; Helsel et al., 2017; Jones et al., 2006; Light et al., 2005; Yun et al., 2004). To explore how ID1 and Notch signaling can allow angioblasts to acquire hemogenic endothelial cell fate, we tested the function of ID1 in EC formation, and its interaction with the Notch signaling pathway. In *mib*^-/-^ mutant embryos, we observed loss of ID1 expression in the endothelium (Figure 9A, B), confirming the results we obtained through scRNA-seq (Figure 8A). Next, we overexpressed NICD in vascular cells using *Cdh5:Gal4; UAS-NICD*, and observed increased expression of ID1 (Figure 9C, D). Together, these results show that Notch signaling regulates the expression of ID1 in the endothelium. Next, to explore potential roles of ID1 in EC fate specification, we validated an ID1 MO (Supplementary Figure 6) and used it to block ID1 gene function. Upon ID1 knockdown, we observe that the expression of endothelial genes *cdh5, notch1b* and *tbx20*, remained unchanged (Figure 9 E-J). However, expression of *runx1*, the key marker of commitment to hemogenic endothelium, was lost (Figure 9K, L). These results show that ID1 is an important regulator of hemogenic cell fate. Next, we were interested in determining the genetic interactions between ID1 and Notch Signaling in the specification of the hemogenic endothelium (Figure 9M-T). Enforced expression of NICD in the vasculature of ID1 morphants embryos failed to rescue *runx1* expression in the hemogenic endothelium (Figure 9M-P). Conversely, enforced expression of ID1 via mRNA injection into *mib*^-/-^ mutant embryos, similarly showed that ID1 gene function is insufficient to rescue loss of Notch signaling in the hemogenic program (Figure 9Q-T). Together, these results demonstrate that ID1 and Notch signaling act in tandem to specify HSPC precursors, likely by maintaining angioblasts in a plastic state to receive the additional cues necessary for acquisition of hemogenic endothelium cell fate.

**Figure 9:**
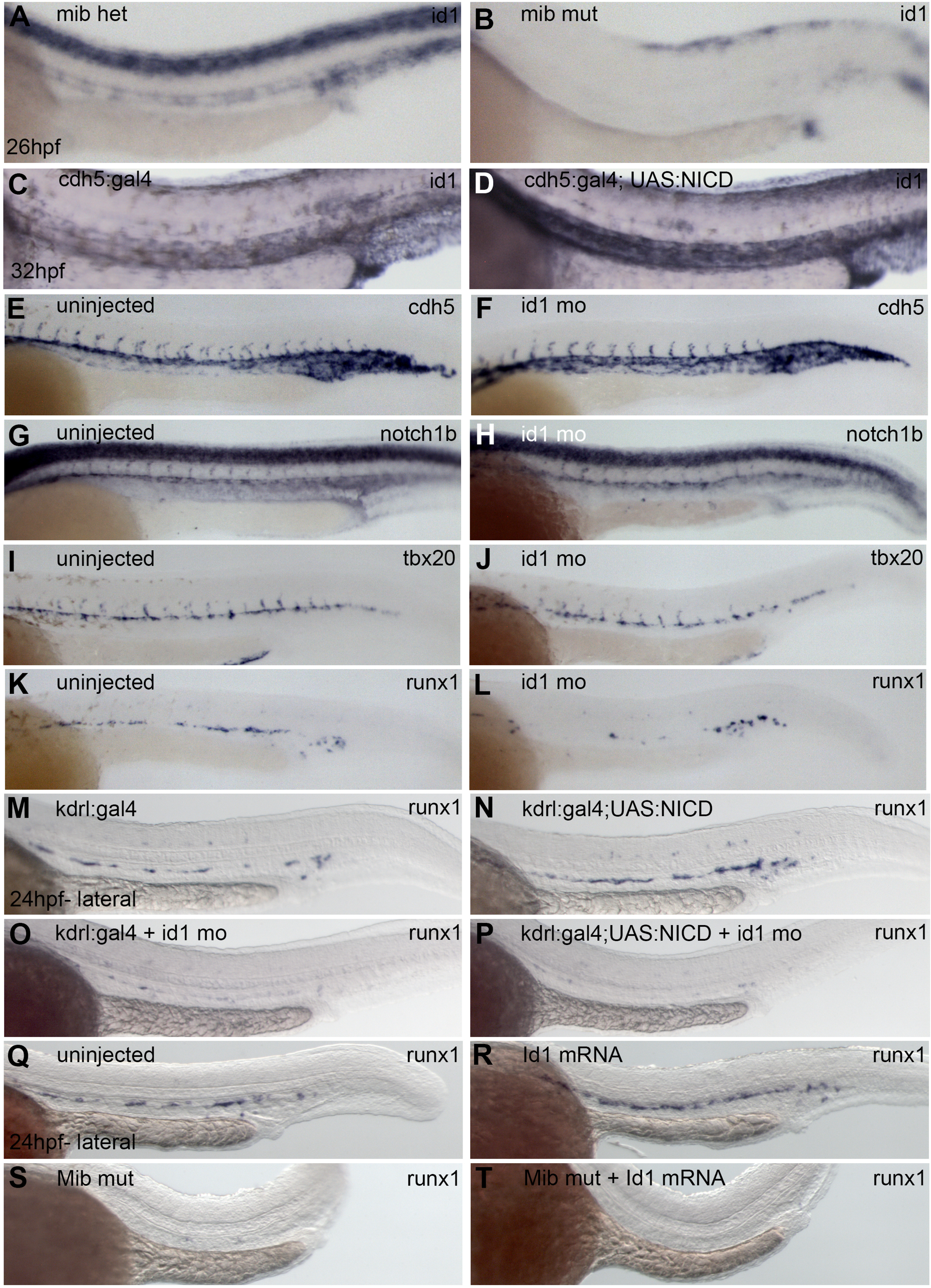
Notch Signaling and ID1 act in tandem for the proper specification of the hemogenic endothelium. **A, B**. mib mutants show a loss of ID1 expression in the neural tube and vasculature region compared to control sibling embryos at 24hpf. **C, D** Conversely, overexpression of Notch signaling in the vasculature region using *Tg(Cdh5:Gal4; UAS:NICD)* results in increased expression of ID1 compared to control sibling embryos. **E-L** Knockdown of ID1 results in normal expression of vascular genes but a substantial reduction of hemogenic marker runx1. **M-P** Vascular overexpression of Notch signaling with *Tg(kdrl:Gal4; UAS:NICD)* is not able to rescue runx1 expression exhibited by embryos injected with ID1 morpholino. **Q-T** Conversely, combined overexpression of ID1 and knockdown of Notch signaling is unable to rescue loss of runx1.

## Discussion

Here, we present an extensive analysis on how a bipotent skeletal muscle progenitor population gives rise to a contingent of endothelial cells that is functionally distinct from endothelial cells specified in the LPM. Through single cell analysis we identify unique molecular signatures sufficient to identify somite derived endothelial cells (SDECs), brain endothelial cells (ECs), kidney ECs and endocardium by 24hpf. Together, these results indicate that cell fate restriction of EC subsets occurs earlier than previously thought. Our analysis of endothelial cells at the single cell level suggests that Notch signaling is dispensable for the acquisition of arterial cell fate but is required for the specification of hemogenic endothelium. In addition, our findings elucidate how Notch signaling works together with inhibitor of differentiation 1 (ID1) to properly maintain a progenitor state that allows for the acquisition of hemogenic endothelial fate. Subsequently, SDECs acts as a secondary molecular input to the hemogenic program that is required for the formation of HSPCs.

Avian and murine studies have previously identified a population of endothelial cells that arise from the somites (Ambler et al., 2001; Esner et al., 2006; Kardon et al., 2002; Mayeuf-Louchart et al., 2014; Mayeuf-Louchart et al., 2016; Pardanaud et al., 1996; Pouget et al., 2006; Wilting et al., 1995; Yvernogeau et al., 2012). In zebrafish, it has been thought that all endothelial and blood cells arise exclusively from the LPM (Childs et al., 2002; Lawson and Weinstein, 2002; Jin et al., 2005; Zhang and Rodaway, 2007; Kohli et al., 2013). Furthermore, angioblasts are thought to be specified as an equipotent population that only upon migration to the midline, undergo progressive cell-fate restriction (Lawson et al., 2001; Kobayashi et al., 2014). To our knowledge there have only been two studies in zebrafish that have shown that the paraxial mesoderm can generate endothelial cells (Martin and Kimelman, 2012; Nguyen et al., 2014). The first noted that presomitic mesoderm could generate ECs that incorporated into caudal blood vessels following inhibition of the Wnt signaling pathway after gastrulation (Martin and Kimelman, 2012). The second noted a rare population of ECs generated in the somite from a central location termed the “endotome” that migrated to the developing dorsal aorta (Nguyen et al., 2014). Each of these studies was based on retrospective analyses, however, making it difficult to ascertain precisely when and where SDECs arise, and their extent of contribution. Use of an *Etv2:GFP* transgene, one of the earliest markers of endothelial cell fate acquisition, allows precise visualization of when and where SDECs emerge from the somites. Furthermore, via use of an endothelial-specific switch transgene, ECs from either the LPM or PM can be permanently and differentially marked to follow their respective contributions to the vasculature. Similar indelible lineage tracing approaches allowed us to rule out any contribution of SDECs to the adult hematopoietic tissue, which is consistent with many previous studies concluding that the adult hematopoietic program is LPM-derived(Henninger et al., 2017; Jin et al., 2007; Murayama et al., 2006). These results are consistent with a previous study from Currie and colleagues that identified a rare population of EC precursors born in the somite that incorporated into the axial vessels but did not appear to generate blood. Their results suggested a central location within the somite that generated ECs, which they termed the “endotome”. In our study, we observed that somite-derived Etv2^+^ ECs arise from the hypaxial dermomyotome, consistent with earlier observations in avian and mouse embryos (Eichmann et al., 1993; Ema et al., 2006; Pouget et al., 2006; Tozer et al., 2007). We observed no ECs from central locations within the somite. Notably, the hypaxial dermomyotome is comprised of bipotent-skeletal muscle progenitors that upon local cues generate ECs or myoblasts. Since these bipotent progenitors are present in the territory where the hypaxial muscle precursors originate (Christ and Ordahl, 1995), we believe a more appropriate term for this somitic compartment would be the ‘Myo-Endotome’.

We observe that Etv2^+^ SDECs emerge starting at the 12-somite stage, concomitant with the epithelization of the somite and the migration of LPM-derived ECs to the midline. Previous work in mouse had suggested that Notch signaling promotes the specification of SDECs at the expense of muscle cell fates (Mayeuf-Louchart et al., 2014). By contrast, we find that Notch signaling is essential for the maintenance of muscle progenitors, as previously shown in mice (Schuster-Gossler et al., 2007). Loss of the Notch signaling components Mib or Notch3 results in a premature depletion of skeletal muscle progenitor markers, and a concomitant increase of differentiated muscle and endothelial cell genes. Therefore, our results show that Notch signaling is required for the maintenance of muscle progenitors and not the cell fate choice between muscle and endothelium. In addition, we find Wnt signaling to be a necessary factor in the regionalization of muscle and endothelial cells within the somites, in agreement with previous work (Borello et al., 2006; Martin and Kimelman, 2012). We find that during homeostasis, few Etv2:GFP^+^ SDECs delaminate from the hypaxial dermomyotome starting from somite number 12. However, combined loss of Notch signaling and Meox1 shows that EC competence within the somite encompasses a much larger territory, extending as far as the median dermomyotome (Supplementary Figure 1). Interestingly, we find that endothelial potential within bipotent skeletal muscle progenitors is NPAS4l-dependent. These results are surprising, since NPAS4l has been shown to be the earliest gene required for EC specification and was thought to be restricted to the LPM starting at the tailbud stage (Reischauer et al., 2016). Furthermore, expression of NPAS4l decreases sharply during early somitogenesis. Together, these results suggest that NPAS4l may provide endothelial competence to an early mesodermal precursor, or may act in a non-cell autonomous manner. Future work should help clarify the exact function of NPAS4l and its activity across mesodermal derivatives.

Our single-cell analyses identified an endothelial population with PM gene signatures that we believe are indicative of SDECs. Our work also classified distinct endothelial clusters at 22-24hpf that exhibit unique molecular signatures that correspond to brain endothelium, kidney endothelium, and endocardium. Interestingly, our results also suggest that SDECs emerge from shared muscle progenitors that arise from the lateral region of the PM that diverges from LPM progenitors by mid-gastrulation (Garcia-Martinez and Schoenwolf, 1992; Schoenwolf et al., 1992; Selleck and Stern, 1991; Psychoyos and Stern, 1996). Future work should help determine if distinct regions within the mesoderm can give rise to functionally divergent subsets of angioblasts. Plasticity of EC development is further illustrated from recent work that surprisingly showed that extraembryonic derived erythomyeloid progenitors can give rise to up to 60% of liver ECs in mice (Plein et al., 2018).

Our single cell analysis of Notch mutant *mib*^*-/-*^; *Etv2:Kaede*^*+*^ ECs provides interesting insights into the role of Notch signaling in early endothelial specification. Using RNA Velocity, our results indicate that endothelial cells at 22hpf are at distinct states along their differentiation hierarchy. Interestingly, when we compare wt *Etv2:Kaede*^*+*^ single cells to those from *mib*^*-/-*^; *Etv2:Kaede*^*+*^ mutants, we find cells further along the differentiation hierarchy in the absence of Notch signaling. Consistent with this finding, we observe transcripts of the arterial gene program to be prematurely expressed. Our results thus highlight a previously underappreciated role for Notch signaling in maintaining angioblasts in a plastic state to regulate their rate of differentiation. Furthermore, our results suggest that early Notch signaling is dispensable for the initiation of the arterial gene program, in agreement with previous studies (Casie Chetty et al., 2017; Wythe et al., 2013), and for the specification of SDECs. Moreover, our single cell analyses mapped Notch mutant Etv2:Kaede^+^ cells to the hemogenic cluster in 22 hpf embryos, which allowed us to investigate the consequences of loss of Notch signaling in hemogenic progenitors. Absence of Notch signaling is associated with a reduction of expression of key hematopoietic genes, including *gfi1aa, runx1*, and *gata2b*. Since we are still able to map mib^-/-^ mutant cells to the hemogenic cluster, this suggests that additional signal(s) acting in concert with Notch are necessary for the correct specification of hemogenic endothelium. Together, our results support the idea that Notch signaling is required for maintaining endothelial progenitors in a plastic state that is required for the proper transition and differentiation towards hemogenic endothelium.

This role for Notch signaling in facilitating the transition to hemogenic endothelium is further illustrated by the role of ID1, one of the most differentially expressed genes between wt and Notch mutant Etv2-kaede^+^ cells. We found Notch signaling to act upstream of ID1 function, and both signaling pathways to be required for the proper specification of hemogenic endothelium. These results suggest that the precursors of hemogenic endothelium need to be maintained in a progenitor state by both Notch and ID1 activity. Once these precursors incorporate into the nascent dorsal aorta, their transition to HSPCs is instructed via HSPCs. Previous work has suggested that the mesenchyme, located directly underneath the dorsal aorta, is a source of BMP ligand necessary for the HSC emergence (Crisan et al., 2015; Durand et al., 2007; Lempereur et al., 2018; Pouget et al., 2014; Wilkinson et al., 2009). More recent studies have also elucidated a role for non-hemogenic ECs in supporting the hemogenic program and in amplifying HSPC number (Butler et al., 2012; Butler et al., 2010; Butler and Rafii, 2012; Gori et al., 2015; Gori et al., 2017; Guo et al., 2017; Hadland et al., 2015; Kim et al., 2014; Kobayashi et al., 2010; Lis et al., 2017; Raynaud et al., 2013; Sandler et al., 2014). Interestingly, several groups have successfully differentiated endothelial precursors into transplantable HSPCs, a process that requires an endothelial ‘niche’ population (Lis et al., 2017; Sandler et al., 2014). Our work indicates that a similar process occurs *in vivo*, where SDECs are required for the induction of hematopoietic cell formation. Future work will elucidate the molecular nature of this requirement and will forward efforts to instruct HSPC fate from human pluripotent precursors.

## Materials and Methods

### Zebrafish husbandry

Wild-type AB* and transgenic TgBAC(kdrl:LOXP-AmCyan-LOXP-ZsYellow) (Zhou et al., 2011) referred to as Kdrl:CSY; Tg(etv2.1:EGFP)zf372 (Veldman and Lin, 2012) referred to as Etv2:GFP; Tg(Tbx6:Gal4FF:GFPnls) (Yabe et al., 2016) referred to as Tbx6:Gal4; TgBAC(etv2:Kaede)ci6 (Kohli et al., 2013) referred to as Etv2:Kaede, Tg(5xUAS-E1B:6xMYC-notch1a) (Scheer et al., 2001) referred to as UAS:NICD; mib^ta52b/ta52b^ (Itoh et al., 2003) referred to as mib^-/-^; Notch3^fh332/fh332^ (Alunni et al., 2013) referred to as Notch3^-/-^; Tg(fli1:DsRed)^um13^(Villefranc et al., 2007) referred to as fli1:DsRed; Tg(Tp1:GFP)^um14^ (Parsons et al., 2009) referred to as Tp1:GFP, Et(ph1bd1:Gal4-mcherry) (Distel et al., 2009) referred to as ph1bd1:Gal4, Tg(cdh5^BAC^:gal4ff)^mu101^ (Bussmann and Schulte-Merker, 2011) referred to as cdh5:Gal4; Tg(Drl:H2Bdendra) (Mosimann et al., 2015) referred to as Drl:H2Bdendra; Tg(Drl:CreERT) (Henninger et al., 2017) referred to as Drl:CreERT; Tg(UAS-Cre)* (Butko et al., 2015) referred to as UAS-Cre. zebrafish embryos and adult fish were raised in a circulating aquarium system (Aquaneering) at 28 °C and maintained in accordance with UCSD Institutional Animal Care and Use Committee (IACUC) guidelines. *Of note, Tg(UAS-CRE) transgene animals were not kept in the background of any Tg(Gal4) transgenic animals, since we noticed it promoted the silencing of the Tg(UAS-CRE) transgene in the next generation.

### WISH

Whole-mount single or double enzymatic in situ hybridization was performed on embryos fixed overnight with 4% paraformaldehyde (PFA) in phosphate buffered saline (PBS). Fixed embryos were washed briefly in PBS and transferred into methanol for storage at –20 °C. Embryos were rehydrated stepwise through methanol in PBS–0.1% Tween 20 (PBT). Rehydrated embryo samples were then incubated with 10 μg ml^−1^ proteinase K in PBT for 5 min for 5 to 10 somite stage (12–15 hpf) embryos and 15 min for 24 to 36 hpf embryos. After proteinase K treatment, samples were washed in PBT and refixed in 4% PFA for 20 min at room temperature. After washes in two changes of PBT, embryos were prehybridized at 65 °C for 1 h in hybridization buffer (50% formamide, 5x SSC, 500 μg ml^−1^ torula tRNA, 50 μg ml^−1^ heparin, 0.1% Tween 20, 9 mM citric acid (pH 6.5)). Samples were then hybridized overnight in hybridization buffer including digoxigenin (DIG)- or fluorescein-labelled RNA probe. After hybridization, experimental samples were washed stepwise at 65 °C for 15 min each in hybridization buffer in 2 × SSC mix (75%, 50%, 25%), followed by two washes with 0.2 × SSC for 30 min each at 65 °C. Further washes were performed at room temperature for 5 min each with 0.2 × SSC in PBT (75%, 50%, 25%). Samples were incubated in PBT with 2% heat-inactivated goat serum and 2 mg ml^−1^ bovine serum albumin (block solution) for 1 h and then incubated overnight at 4 °C in block solution with diluted DIG-antibodies (1:5,000) conjugated with alkaline phosphatase (AP) (Roche). To visualize WISH signal, samples were washed three times in AP reaction buffer (100 mM Tris, pH 9.5, 50 mM MgCl_2_, 100 mM NaCl, and 0.1% Tween 20) for 5 min each and then incubated in the AP reaction buffer with NBT/BCIP substrate (Roche).

For two-color double FISH, embryos were blocked in maleic acid buffer (MAB; 150 mM maleic acid, 100 mM NaCl, pH 7.5) with 2% Roche blocking reagent (MABB) for 1 h at room temperature, after hybridizing at 65 °C with probes as described above. Embryos were incubated overnight at 4 °C in MABB with anti-fluorescein POD (Roche) at a 1:500 dilution. After four washes in MAB for 20 min each followed by washes in PBS at room temperature, embryo samples were incubated in TSA Plus Fluorescein Solution (Perkin Elmer) for 1 h. Embryos were washed 10 min each in methanol in PBS (25%, 50%, 75%, 100%). Embryos were incubated in 1% H_2_O_2_ in methanol for 30 min at room temperature and washed for 10 min each in methanol in PBS (75%, 50% 25%) and 10 min in PBS. After blocking for 1 h in MABB, embryos were incubated overnight at 4 °C in MABB with anti-DIG POD (Roche) at a 1:1,000 dilution. Samples were washed and incubated in TSA Plus CY3 solution (Perkin Elmer) as described above. Embryos were washed three times for 10 min each in PBT and refixed in 4% PFA after the staining was complete. Antisense RNA probes for the following genes were prepared using probes containing DIG or fluorescein labelled UTP: *etv2, meox1, tcf15, pax3a, myoD*,

### Immunofluorescence and microscopy

For immunofluorescence staining for Myc in *hsp70:gal4*; *UAS:NICD-myc* zebrafish embryos after WISH, 100% acetone was added to the embryos (7 min at –20 °C) instead of proteinase K during the WISH procedure described above. WISH samples of embryos were blocked in MABB for 1 h and incubated overnight in MABB with anti-Myc monoclonal 9E10 antibodies at 1:100 (Covance). After four washes in MABB for 30 min each, embryos were incubated in AlexaFuor 488 donkey anti-mouse IgG secondary antibodies (ThermoFisher) at 1:500 in MABB. After staining, embryos were washed in MABB for four times for 30 min each. Live transgenic embryos and flat-mount or whole-mount two-colour double FISH samples were imaged using confocal microscopy (SP5 or SP8 Leica). Embryos were embedded and sectioned as previously described in Kobayashi et al., 2014.

### Pharmacological treatment

IWR (Tocris) was dissolved in DMSO at a concentration of 10 mM. Zebrafish AB* embryos were incubated in 10μM IWR solution from 11 to 15 hpf followed by fixation with 4% PFA or snap frozen for RNA collection.

### Microinjections of morpholinos, mRNA and plasmids

Antisense morpholinos (MOs; Gene Tools, LLC) were diluted as 1- or 3-mM stock in H_2_O. Meox1-MO, Mib-MO, and ID1-MO were injected at 1-to 2-cell stage of development. The sequence of translation-blocking targeted ID1-MO1 is 5’- -3’. Full-length *ID1* and mOrange-CAAX mRNAs were synthesized from linearized pCS2+ *ID1* or mOrange-CAAX with the mMessege mMaching kit (Ambion). *ID1* and mOrange-CAAX mRNA were injected with 100 pg into 1-to 2-cell stage embryos. 25pg of Mylz2:Etv2 construct was coinjected with 50pg of Transposase mRNA. All microinjections were performed with the indicated concentration of RNA or MO in a volume of 1 nl using a PM 1,000 cell microinjector (MDI).

### FACS

*Tg(Etv2:kaede;mib*^*-/-*^) and Tg(E*tv2:kaede;mib*^*+/+*^*), Tg(* E*tv2:kaede;mib*^*+/-*^), were separated into two distinct pools at 22 hpf based on their head phenotype as previously described (Bingham et al.). Tg(Fli1:dsRed;Tp1GFP) and Tg(Draculin:Dendra) were separately dissociated in PBS supplemented with 10%FBS by homogenizing with sterile plastic pestle or pipette. Dissociated cells were filtered through a 35-μm nylon cell strainer (Falcon 2340) and then rinsed with PBS with 10% FBS. Propidium iodide (Sigma) was added (1 μg ml^−1^) to exclude dead cells and debris. FACS was performed based on GFP, Dendra, and DsRed fluorescence with a FACS Aria II flow cytometer (Beckton Dickinson).

### Quantitative real-time RT-PCR

Total RNA was collected from whole embryos (∼20 embryos) using TRIzol reagent (Ambion, Life Technologies) and isolated from Notch3^-/-^ mutant embryos or control siblings at 48hpf; Cloche^-/-^ mutant embryos or control siblings at 48hpf; or IWP2 treated and control embryos at 15hpf with the RNeasy kit (Qiagen). cDNA was generated from total RNA with iScript cDNA synthesis kit (Bio Rad). The following primers were used for cDNA quantification: ef1α (forward, 5’- GAGAAGTTCGAGAAGGAAGC -3’; reverse, 5’- CGTAGTATTTGCTGGTCTCG -3’), etv2 (forward, 5’- CGAGGTTCTGGTAGGTTTGAG - 3’; reverse, 5’- GCACAAAGGTCATGTTCTCAC -3’), fli1a (forward, 5’- CGTCAAGCGAGAGTATGACC -3’; reverse, 5’- AGTTCATCTGAGACGCTTCG -3’), myod1 (forward, 5’- GAAGACGGAACAGCTATGAC -3’; reverse, 5’- GGAGTCTCTGTGGAAATTCG -3’), myf5 (forward, 5’- CCAGACAGTCCAAACAACAGACC -3’; reverse, 5’- TGAGCAAGCAGTGTGAGTAAGCG -3’), pax3a (forward, 5’- ATTCCTTGGAGGTCTCTACG -3’; reverse, 5’- CTACTATCTTGTGGCGGATG -3’), pax7b (forward, 5’- CAGTATTGACGGCATTCTGGGAG -3’; reverse, 5’- TCTCTGCTTTCTCTTGAGCGGC -3’), myf5 (forward, 5’- GAATAGCTACAACTTTGACG -3’; reverse, 5’- GTAAACTGGTCTGTTGTTTG -3’), myog (forward, 5’- GTGGACAGCATAACGGGAACAG -3’; reverse, 5’- TCTGAAGGTAACGGTGAGTCGG-3’).

### Single-cell RNA sample preparation

After FACS, total cell concentration and viability were ascertained using a TC20 Automated Cell Counter (Bio-Rad). Samples were then resuspended in 1XPBS with 10% BSA at a concentration between 800-3000 per ml. Samples were loaded on the 10X Chromium system and processed as per manufacturer’s instructions (10X Genomics). Single cell libraries were prepared as per the manufacturer’s instructions using the Single Cell 3’ Reagent Kit v2 (10X Genomics). Single cell RNA-seq libraries and barcode amplicons were sequenced on an Illumina HiSeq platform.

### Single-cell RNA sequencing analysis

The Chromium 3’ sequencing libraries were generated using Chromium Single Cell 3’ Chip kit v3 and sequenced with (actually, I don’t know:(what instrument was used?). The Ilumina FASTQ files were used to generate filtered matrices using CellRanger (10X Genomics) with default parameters and imported into R for exploration and statistical analysis using a Seurat package (La Manno et al., 2018). Counts were normalized according to total expression, multiplied by a scale factor (10,000), and log transformed. For cell cluster identification and visualization, gene expression values were also scaled according to highly variable genes after controlling for unwanted variation generated by sample identity. Cell clusters were identified based on UMAP of first 14 principal components of PCA using Seurat’s method, Find Clusters, with a original Louvain algorithm and resolution parameter value 0.5. To find cluster marker genes, Seurat’s method, FindAllMarkers. Only genes exhibiting significant (adjusted p-value < 0.05) a minimal average absolute log2-fold change of 0.2 between each of the clusters and the rest of the dataset were considered as differentially expressed. To merge individual datasets and to remove batch effects, Seurat v3 Integration and Label Transfer standard workflow (Stuart et al., 2019).

### Plasmid Construction

Plasmids Expression constructs were generated by standard means using PCR from cDNA libraries generated from zebrafish larvae at 24 hpf, these were cloned into pCS2+, downstream of a SP6 promoter. Previously described Mylz2 promoter (Ju et al., 2003) was cloned upstream of Etv2 CDS flanked by Tol2 sites.

## Replicates

All experiments for the assessment of phenotype and expression patterns were replicated in at least two independent experiments. Embryos were collected from independent crosses, and experimental processing (staining or injections) were carried out on independent occasions. Exceptions to this include the analysis of the kidney hematopoietic cells by flow from Tbx6:Gal4; UAS-CRE; A2BD and the imaging of Tbx6:Gal4; UAS-CRE; Kdrl-CSY where many embryos where processed, and the corresponding n are reported in the associated figure legends. The analysis of qRTPCR where 3 independent experiments were performed per condition; and the single-cell sequencing experiments, where one sequencing run was performed per time-point from a pool of at least 100 embryos within the corresponding transgenic background.

## Acknowledgements

We thank members of the Traver laboratory for helpful discussions, the UCSD Institute for Genomic Medicine sequencing core for their support on the scRNA-seq sample preparation and sequencing, and Stephanie Grainger for discussions and careful editing of the manuscript.

## Author Contributions

PS-H and CP conceived, designed and conducted experiments and analysis, and wrote the manuscript. OS conducted experiments and analysis. DT supervised experiments and edited the manuscript.

## Funding

This work was supported by the National Institute of Health [R01-DK074482 to D.T. and P.S.-H.]; National Institute of Health [T32-HL086344 to P.S.-H.]; OS was funded by American Heart Association [19POST34380328].

---

**Supplementary Figure 1:**
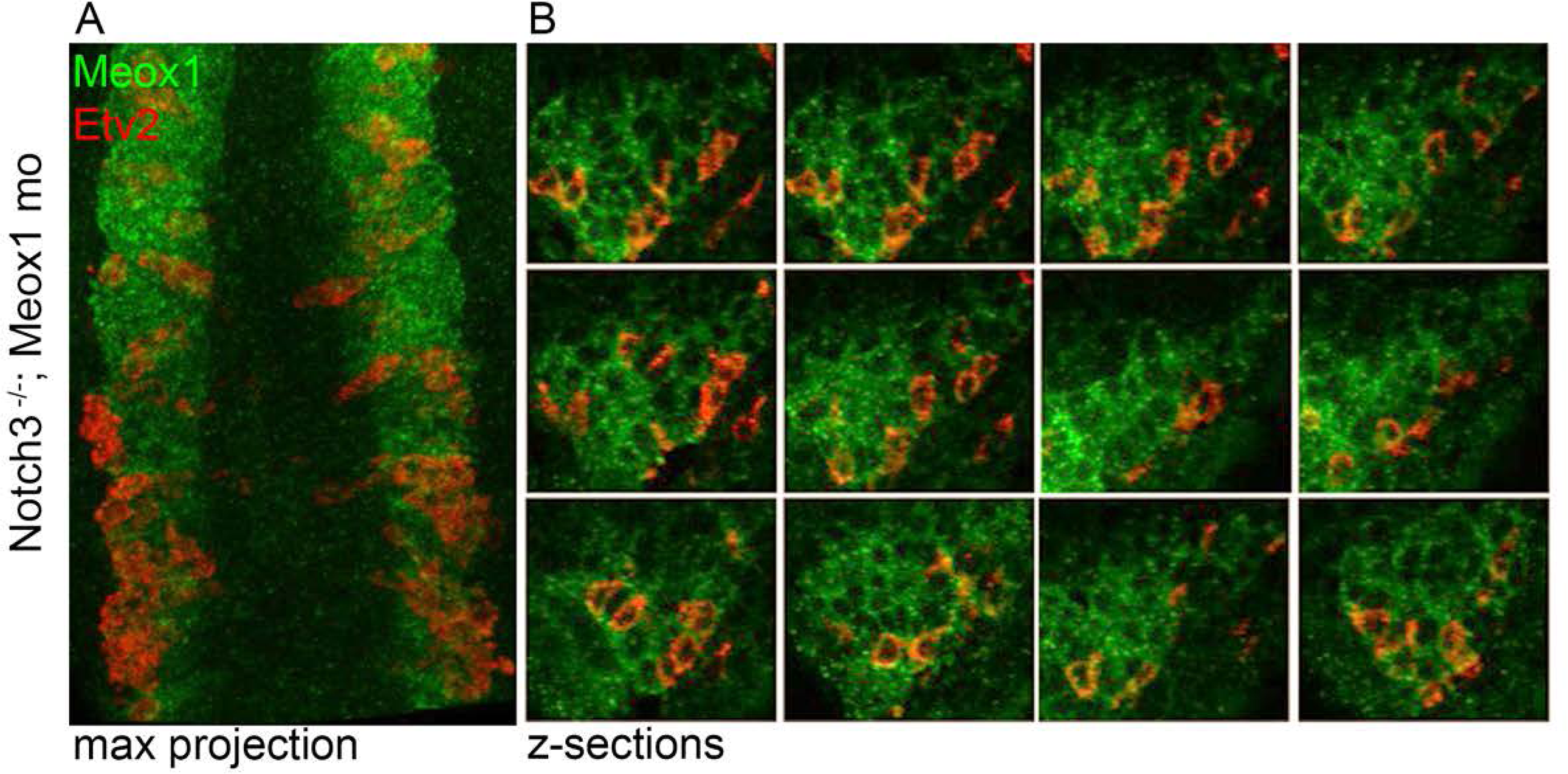
Bipotent muscle progenitor cells contain endothelial potential that can span deep into the dermomyotome compartment. **A**. Max projection of 12hpf Notch3 mutant embryos injected with meox1 morpholino shows the extent of endothelial potential within the somite compartment (double positive *meox1* (green) and *etv2* (red) cells). **B**. Z-sections of a dermomyotome compartment, that shows the extent of double positive cells.

**Supplementary Figure 2:**
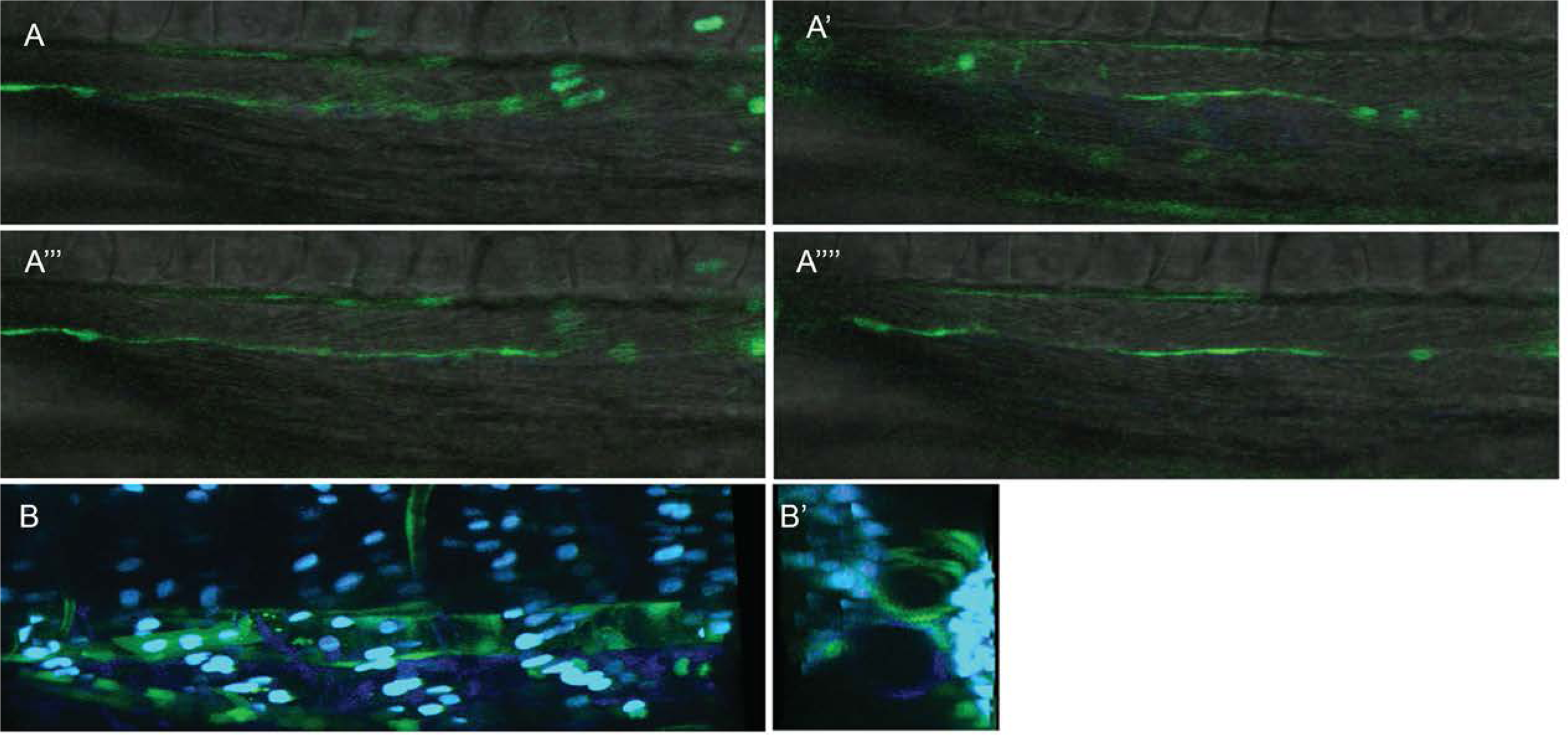
Paraxial mesoderm derived endothelial cells contribute to the dorsal aorta in zebrafish. **A, B**. Cross section and 3D reconstruction shows contribution of *Tg(Tbx6:Gal4;UAS:CRE, Kdrl:CSY)* SDEC YFP^+^ cells to the anterior vasculature region of the dorsal aorta. A-A”” Shows different z-sections of the image. B, B’ shows 3D reconstruction.

**Supplementary Figure 3:**
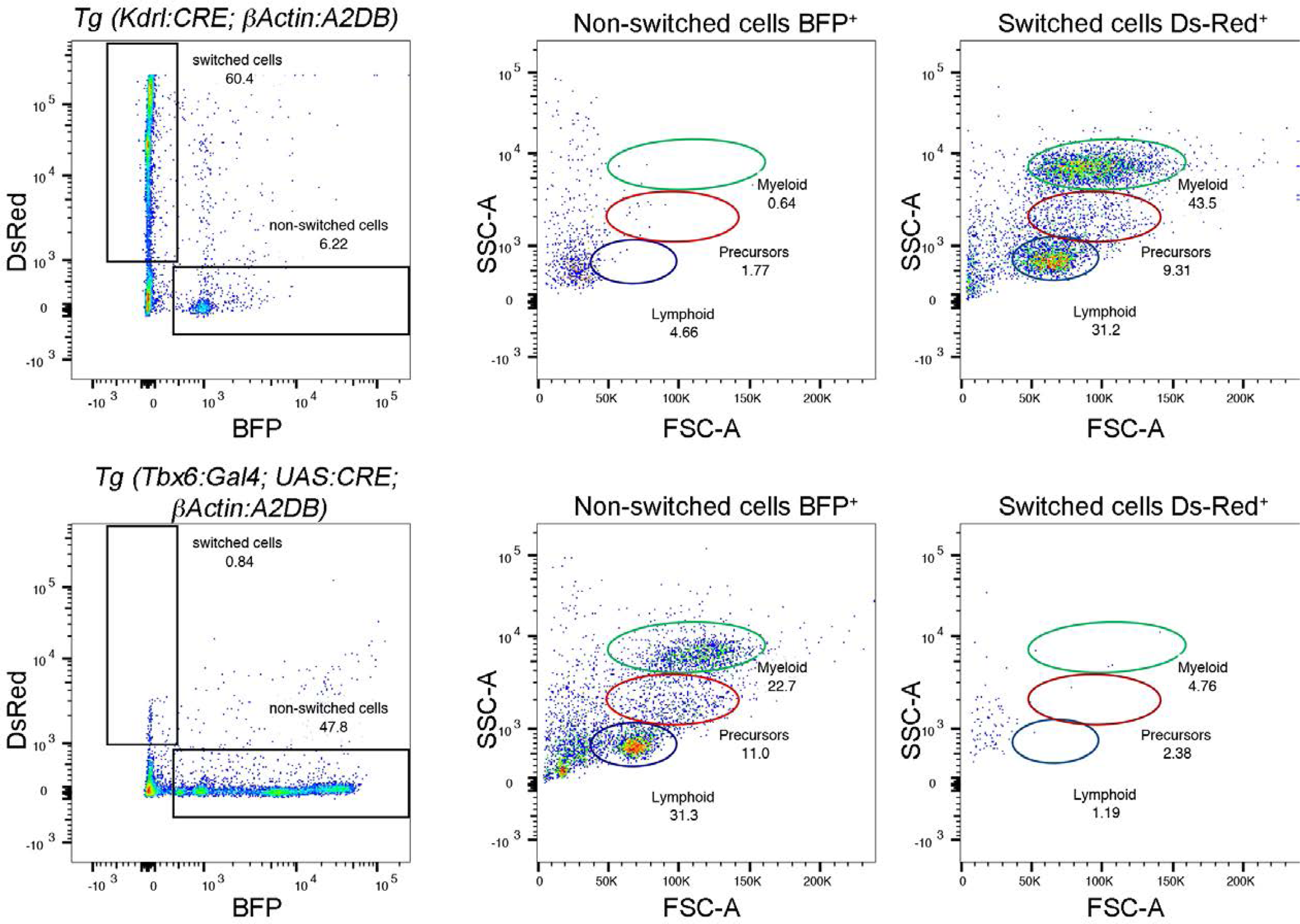
The paraxial mesoderm does not generate HSPCs. *Tg(Tbx6:Gal4; UAS:CRE; beta-actin:A2BD)* and *Tg(Kdrl:CRE; beta-actin:A2BD*) adult kidney marrows were analyzed by flow cytometry. Top row illustrates that the hematopoietic lineages of *Tg(Kdrl:CRE; beta-actin:A2BD)* originate from switched DsRed^+^ cells whereas in the *Tg(Tbx6:Gal4; UAS:CRE; beta-actin:A2BD)* BFP^+^ non-switched cells are the source of the hematopoietic system.

**Supplementary Figure 4:**
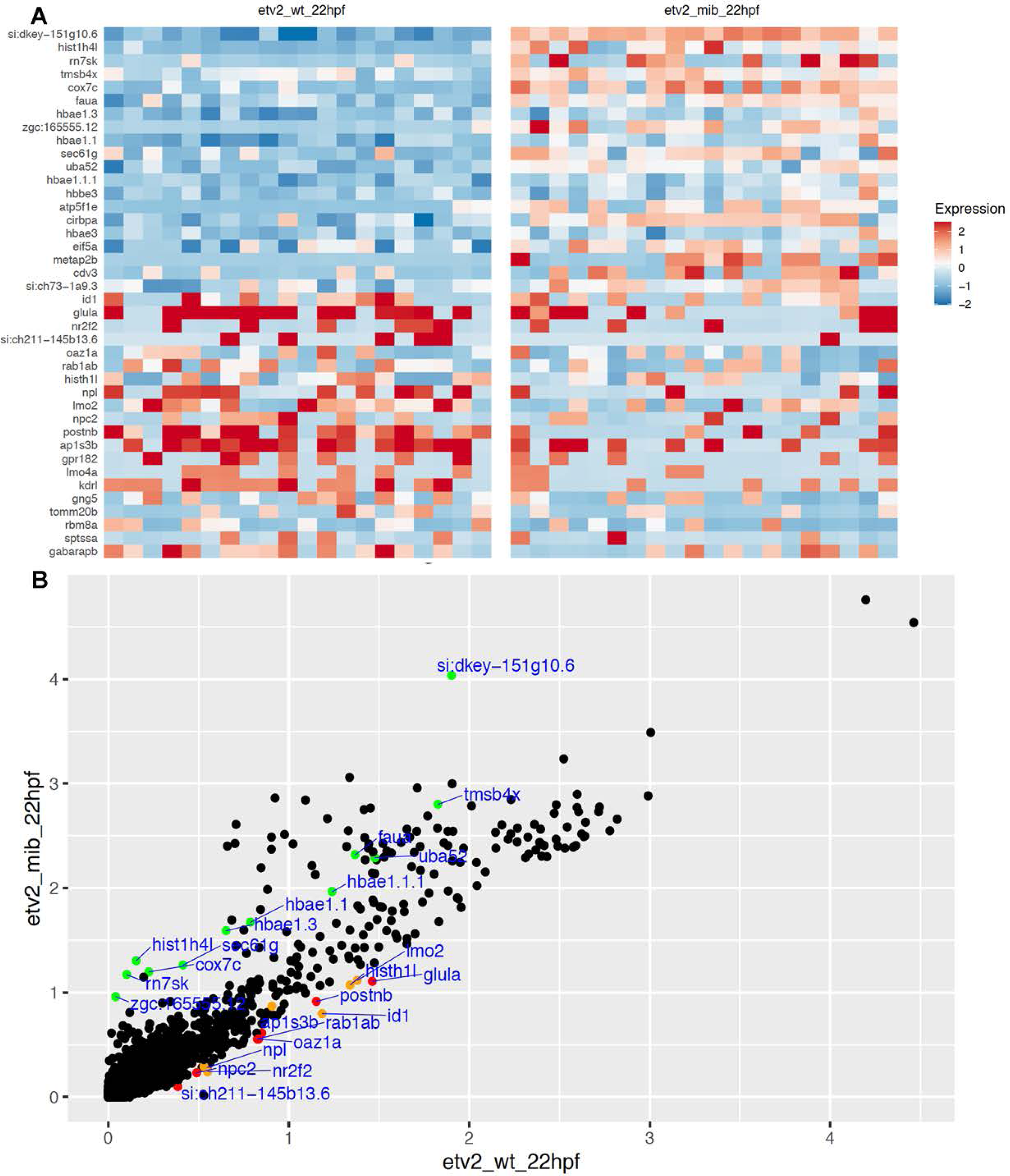
mib^-/-^ single cell analysis in hemogenic endothelial cluster shows that loss of Notch signaling results in reduced expression of genes important for hematopoietic cell formation. **A**. Heatmap of genes specific for the Pre-HSPCs and hemogenic cluster showing the 20 most upregulated and downregulated genes in wild type Etv2:GFP^+^ and mib^-/-^ Etv2:GFP^+^ cells isolated at 22hpf. **B**. Scatter plot of differentially expressed genes between mib^-/-^mutant (y-axis) and control cells (x-axis). Green dots are genes upregulated in Notch mutants. Red dots are genes which expression is decreased in Notch mutants. Genes important for hematopoiesis are represented in orange.

**Supplementary Figure 5:**
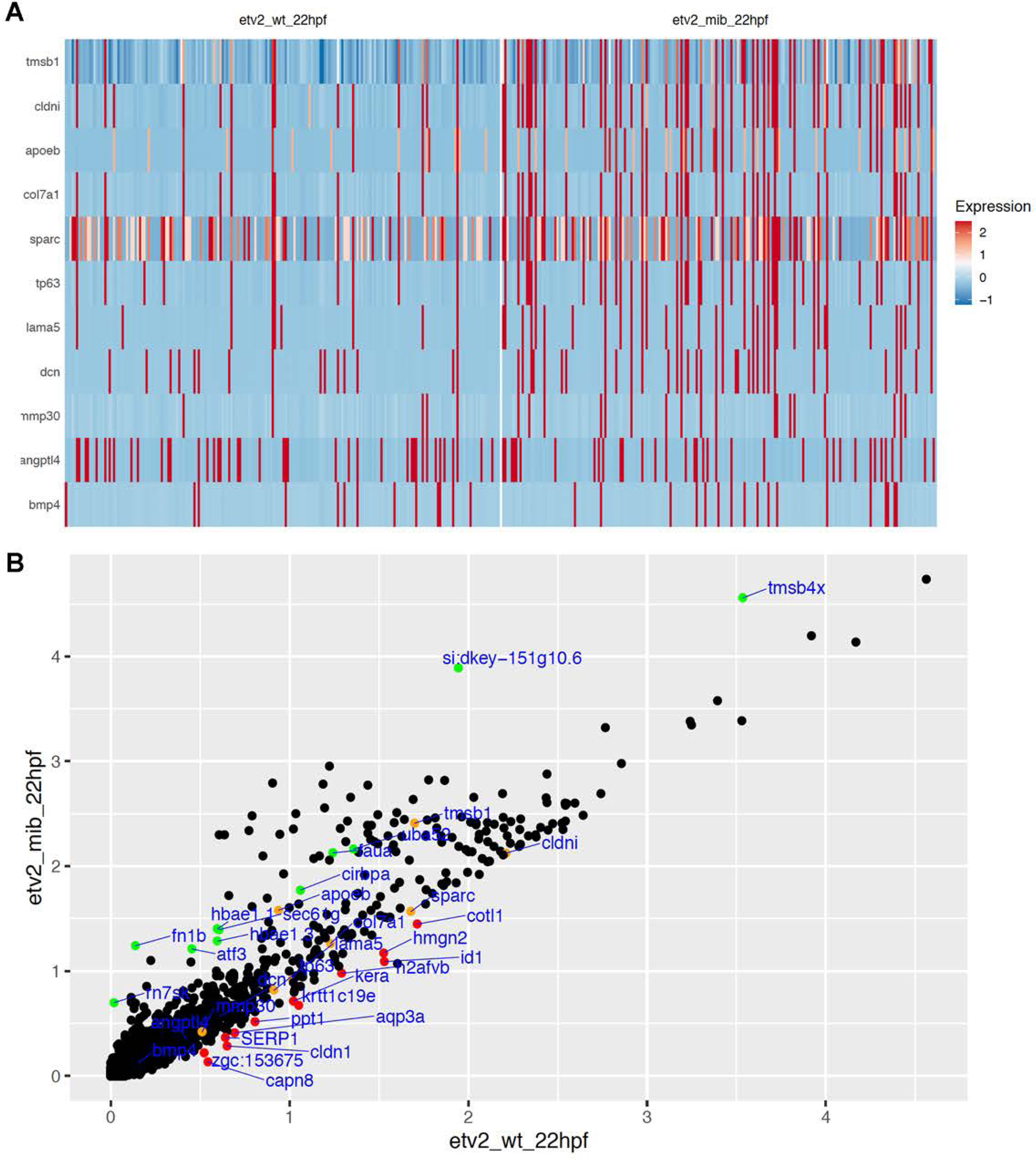
Single cell analysis in mib^-/-^ in SDEC cluster shows that loss of Notch signaling does not affect the specification of SDECs. **A**. Heatmap of genes expressed in SDECs. Genetic ablation of Notch signaling has little impact on SDECs genes. **B**. Scatter plot of differentially expressed genes between mib mutant (y-axis) and control cells (x-axis). Green dots are genes upregulated in Notch mutants. Red dots are genes which expression is decreased in Notch mutants. Genes highly expressed in SDECs are represented in orange.

**Supplementary Figure 6:**
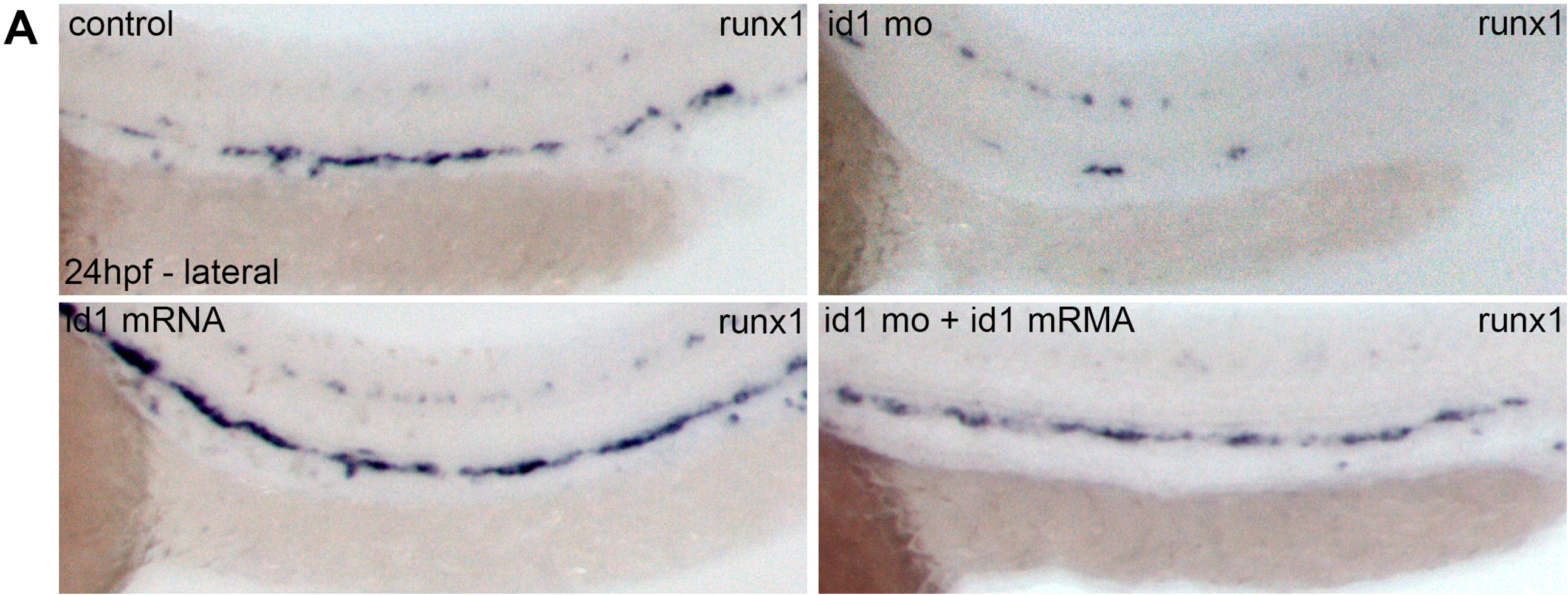
ID1 morpholino validation. Knockdown of Id1 splice morpholino reduced runx1 expression. Conversely, increased dose of Id1 achieved by mRNA injection, upregulated runx1 expression in the dorsal aorta. Combined morpholino knockdown and enforced expression of Id1 restores runx1 expression to a level similar to what is observed in control sibling embryos.

**Supplementary MovieS1: Time-lapse imaging of *Tg(Etv2:GFP, Ph1db1:Gal4-UAS:mCherry)* embryos imaged between 12-16ss**. Lateral view of a transgenic embryo. LPM cells are on the right and migrate under the somites. SDECs arise from the third and the fourth somites while most of the LPM cells have already ingress underneath the somites.

**Supplementary MovieS2: Time-lapse imaging of IWP-L6 treated embryos in the background *Tg(Etv2:GFP)* injected with mOrange2:CAAX mRNA and imaged dorsally between 10-14ss**. Embryos treated with the Wnt inhibitor IWP-L6 between 2ss-14ss observe the ectopic formation of Etv2:GFP+ cells in the central part of the somite

